# The type of DNA damage response after Decitabine treatment depends on the level of DNMT activity

**DOI:** 10.1101/2023.09.02.556017

**Authors:** Tina Aumer, Linda Bergmayr, Stephanie Kartika, Theodor Zeng, Qingyi Ge, Grazia Giorgio, Maike Däther, Alexander J. Hess, Stylianos Michalakis, Franziska R. Traube

## Abstract

Decitabine and Azacytidine are considered as epigenetic drugs that induce DNA-methyltransferase (DNMT)-DNA crosslinks, resulting in DNA-hypomethylation and -damage. Although they are applied against myeloid cancers, important aspects of their mode of action remain unknown, which highly limits their clinical potential. Using a combinatorial approach, we reveal that the efficacy profile of both compounds primarily depends on the level of induced DNA-damage. Under low DNMT-activity, only Decitabine has a substantial impact. Conversely, when DNMT-activity is high, toxicity and cellular response to both compounds are dramatically increased, but do not primarily depend on DNA-hypomethylation or RNA-associated processes, contradicting an RNA-dependent effect of Azacytidine. By applying spatial proteomics, we show that Decitabine induces a strictly DNMT-dependent multifaceted DNA-damage response based on chromatin-recruitment of various repair-associated proteins. The choice of DNA-repair pathway herby depends on the severity of Decitabine-induced DNA-lesions. While mismatch (MMR) and base-excision DNA repair (BER) as well as RAD50- dependent DNA double-strand break repair are always activated in response to Decitabine, Fanconi anemia-dependent DNA-repair combined with homologous recombination is only activated when DNMT-activity is moderate. In contrast, high DNMT-activity and therefore immense replication stress, induce DNA repair by non-homologous and alternative end-joining.

## INTRODUCTION

5-Aza-2’-deoxycytidine (Decitabine, AzadC) and 5-azacytidine (Azacytidine, AzaC) are cytosine analogues that covalently trap DNA-methyltransferases (DNMTs) and therefore belong to the compound class of hypomethylating agents (HMAs) (1,2). Both compounds are applied in the clinic against myelodysplastic syndrome (MDS) and acute myeloid leukaemia (AML), which have otherwise very limited treatment options (3,4). It has been reported that AzadC and AzaC have potentially beneficial effects for therapy of solid tumours as well, but it is not understood yet why solid tumours do not equally respond towards AzadC or AzaC exposure as hematopoietic malignancies (5–8). AzaC and AzadC feature a multi-mode of action (multi-MoA) by addressing epigenetic and DNA damage processes and additionally for AzaC, also RNA-dependent processes (Figure 1A) (1,9). Both compounds are taken up into cells via nucleoside transporters in the plasma membrane, however with different transportability profiles in respect to the different nucleoside transporter types (10). After uptake, 80 – 90% of AzaC is incorporated into RNA, where it inhibits the tRNA (cytosine(38)-C(5))-methyltransferase (*TRDMT1*, *DNMT2*), suggesting an RNA-dependent effect, which is, however, not well characterized yet (9,11,12). The remaining 10 – 20% of AzaC are converted on the diphosphate level to the respective AzadC analogue and the metabolic pathways of AzaC and AzadC unite (Figure 1A). The AzadC-triphosphate can be subsequently used by DNA-polymerases for genomic incorporation of AzadC during S-phase instead of 2’-deoxycytidine (dC). After genomic incorporation, AzadC is recognized by DNMTs as dC but due to the CH-N replacement at the pyrimidine ring, the enzyme cannot be released anymore resulting in permanent crosslinks between the protein and the 5-aza-cytosine nucleobase (1) (Figure 1A). On the epigenome level, DNMT inhibition leads to a global loss of the epigenetic mark 5-methyl-2’-deoxycytidine (mdC), a key player of epigenetic modulation of gene expression (13). This feature can be highly beneficial for tumour therapy as many cancer types and in particular AML subtypes have silenced tumour suppressor genes by hypermethylation of the respective promoter regions (14–17). On the DNA damage level, AzadC can potentially create various types of DNA lesions (Figure 1A). First, AzadC is a non-canonical base, which can rapidly undergo hydrolyzation (18), resulting in DNA lesions by mismatches and abasic sites. However, the most severe form of DNA-damage after exposure to AzadC, which also determines the mutagenicity of AzadC, are the DNMT-DNA crosslinks (19–21). To maintain a level of genomic integrity that is necessary to survive, cancer cells often have a very efficient, but less precise DNA repair machinery (22). This feature not only gives them a survival advantage to deal with naturally occurring DNA lesions, but also to deal with DNA lesions introduced by DNA damaging reagents (23). One clinically important example for chemoresistance by proficient DNA repair is the on-target resistance of many aggressive tumours against cisplatin, which is applied as a cytostatic agent against solid tumours (24). To understand the MoA and the associated repair mechanisms of a DNA-damaging reagents is therefore of utmost importance to improve therapy efficiency in the clinic. Previous studies showed that non-homologous end-joining (NHEJ), homologous recombination (HR) via the Fanconi anemia (FA) pathway and also Poly(ADP-ribose) polymerase 1 (PARP1)-dependent DNA repair are involved in the repair of AzadC-induced DNA lesions (25–27). However, it has not been investigated in a holistic and systematic manner yet which DNA damage responses (DDRs) are activated as a cellular response towards AzadC or AzaC under different cellular prerequisites. This information, however, is pivotal to understand and break resistance mechanisms during AzadC- or AzaC-based cancer therapy.

**Figure 1:**
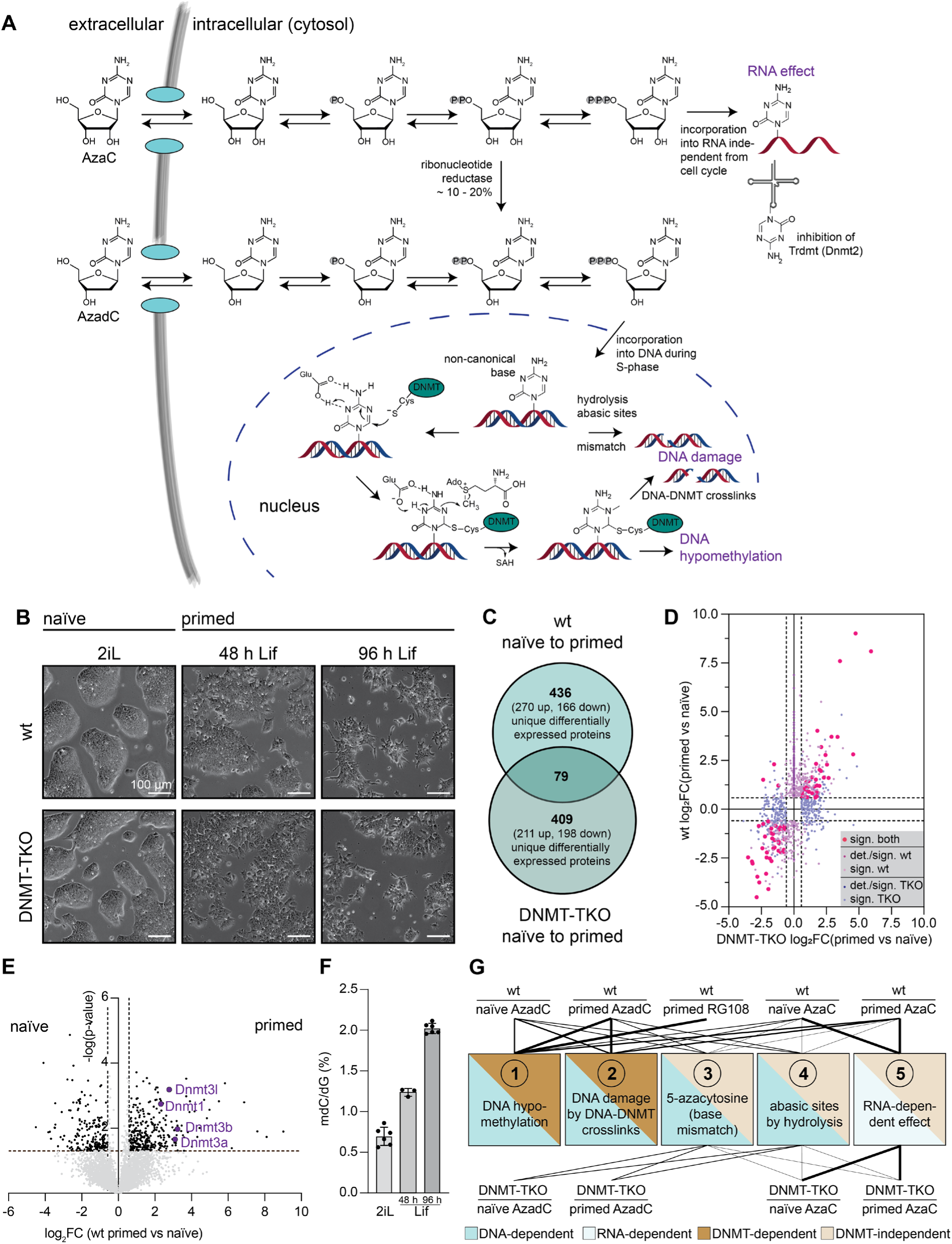
Mouse embryonic stem cells (mESCs) as a model system to study the different modes of action (MoAs) of AzadC and AzaC. A) Schematic representation of the metabolization of AzadC and AzaC in the cell. B) Morphological changes as indicated by representative brightfield microscopy of wt and DNMT-TKO mESCs in the transition from the naïve state (cells cultured in 2iL) to the primed state (cells cultured in Lif). C) Venn diagram of the number of proteins in the wt and the DNMT-TKO that showed significant expression level changes (log_2_FC ≥ |0.58496| and p-value < 0.05) from the naïve to the 96 h primed state. 79 proteins were significantly differentially expressed in both genotypes. D) Correlation plot showing the log_2_FC of the differentially expressed proteins of C). Proteins that were differentially expressed in both genotypes are displayed in magenta, proteins with significant expression level changes that were only detected in the wt are showed in deep purple. Proteins that were detected in both genotypes but only significant in the wt are light purple. Proteins with significant expression level changes that were only detected in the DNMT-TKO are showed in dark blue and proteins that were detected in both genotypes but only significant in the DNMT-TKO are light blue. E) Volcano plot showing the protein expression changes (log_2_FC) and the consistency of the change (-log(p-value)) between the naïve and the primed state in the wt. Proteins with higher expression in the 96 h primed state are on the right side, proteins with higher expression in the naïve state are shown on the left side. Dnmt1, Dnmt3a, Dnmt3b and the DNMT3-regulatory unit Dnmt3I are displayed in purple. Proteins with significant expression changes (|log_2_FC| > 0.58496, −log(p-value) > 1.3 (≙ p-value < 0.05)) are shown in black, rest is shown in grey. F) Quantification of absolute 5mdC in the DNA by triple-quadrupole mass spectrometry, normalized to the amount of dG. G) Diagram of the five possible different MoAs of 5-azacytosine-based compounds and how much they are potentially affected using either wt or DNMT-TKO under different treatment conditions. The more intense the line is, the more influence of the MoA on the cell is expected under the different conditions.

The multi-MoA profiles of AzadC and AzaC provide unique drug profiles in comparison to other chemotherapeutic agents that target either genomic integrity or other cellular features like epigenome patterns. However, to date it is unclear how much the individual MoAs contribute to the efficacy profile of AzadC and AzaC and reliable biomarkers to predict a patient’s response to both, exclusively one or none of the two compounds are still missing. To investigate in detail how much the different MoAs of AzadC and AzaC contribute to their efficacy profile, we chose mouse embryonic stem cells (mESCs) as a model system because their special cellular features allowed us to distinguish the different MoAs from each other (Table 1). In contrast to other cell types, mESCs in a non-committed state can tolerate ablation of Dnmt1, Dnmt3a and Dnmt3b (DNMT-TKO) (Figure S1A, B), resulting in global DNA demethylation (Figure S1C) without showing deleterious developmental and cellular defects (28). Furthermore, mESCs also show sustained proliferative signalling and replicative immortality like cancer cells, but do not feature genomic instability. Importantly, they have a fully functional DDR, which makes them exceptionally suitable to study potential proliferation inhibitory and cell-death inducing effects of AzadC and AzaC as well as involved DNA repair processes. Last, mESCs can be cultured in a naïve (2iL culture conditions) or in a primed state (Lif culture conditions), which represents two pluripotent but distinct developmental stages, where the naïve state is characterized by a very low, but the primed state by a very high DNMT activity (29,30).

**Table 1:** Modes of action (MoA) of HMAs and how they can be distinguished from each other. RG108 is a non-nucleoside based DNMT inhibitor.

| Mode of action (MoA) | Conditions, in which the MoA is present | Conditions, in which the MoA is the only or the dominant MoA compared to other conditions |
| --- | --- | --- |
| 1. DNA hypomethylation | wt with AzadC, AzaC and RG108 | Only MoA in RG108-treated wt cells |
| 2. DNA damage by DNA-DNMT crosslinks | wt with AzadC and AzaC | Dominant MoA in AzadC-treated wt cells when effects of MoA1 are subtracted |
| 3. & 4. DNA lesions by mismatch and formation of abasic sites | wt and DNMT-TKO with AzadC and to a much lesser extent (10 – 20%) with AzaC | Only MoA in AzadC-treated DNMT-TKO cells |
| 5. RNA-dependent effect | wt and DNMT-TKO with AzaC | Dominant MoA in AzaC-treated DNMT-TKO cells |

To dissect the individual contribution of the different MoAs and gain a holistic picture of associated DNA repair mechanisms, we started with AzadC treatment of the wt and the DNMT-TKO in the naïve state to distinguish DNMT-dependent effects from other DNA-based effects. Next, we studied the effects of AzadC on both genotypes in the primed state when DNMT-activity is high compared to the naïve state. Last, we had a closer look on the effects of AzaC in the naïve and the primed state in both genotypes and also investigated the effects of DNMT-inhibition without creating DNA-damage using the previously reported non-nucleoside DNMT inhibitor RG108 (31,32).

## MATERIAL AND METHODS

If not indicated otherwise, Milli-Q grade water was used for all experiments and room temperature (RT) refers to a temperature between 20 °C and 22 °C.

### Reagents and Cell Lines

#### Chemicals

All chemicals that were used in this study are listed in Table 2. If not indicated otherwise, chemicals were used without further purification and stored according to the available product sheet.

**Table 2:**
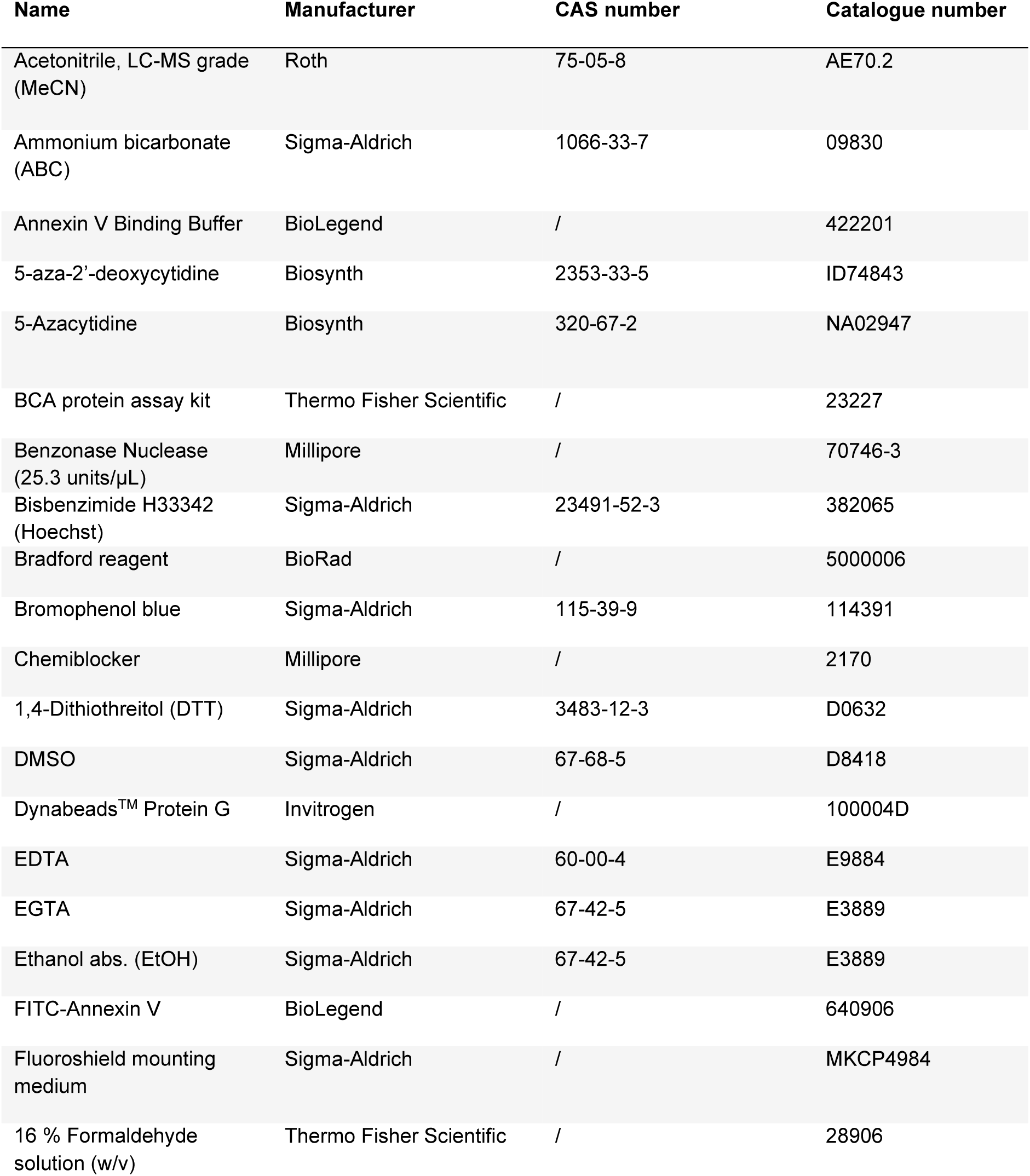
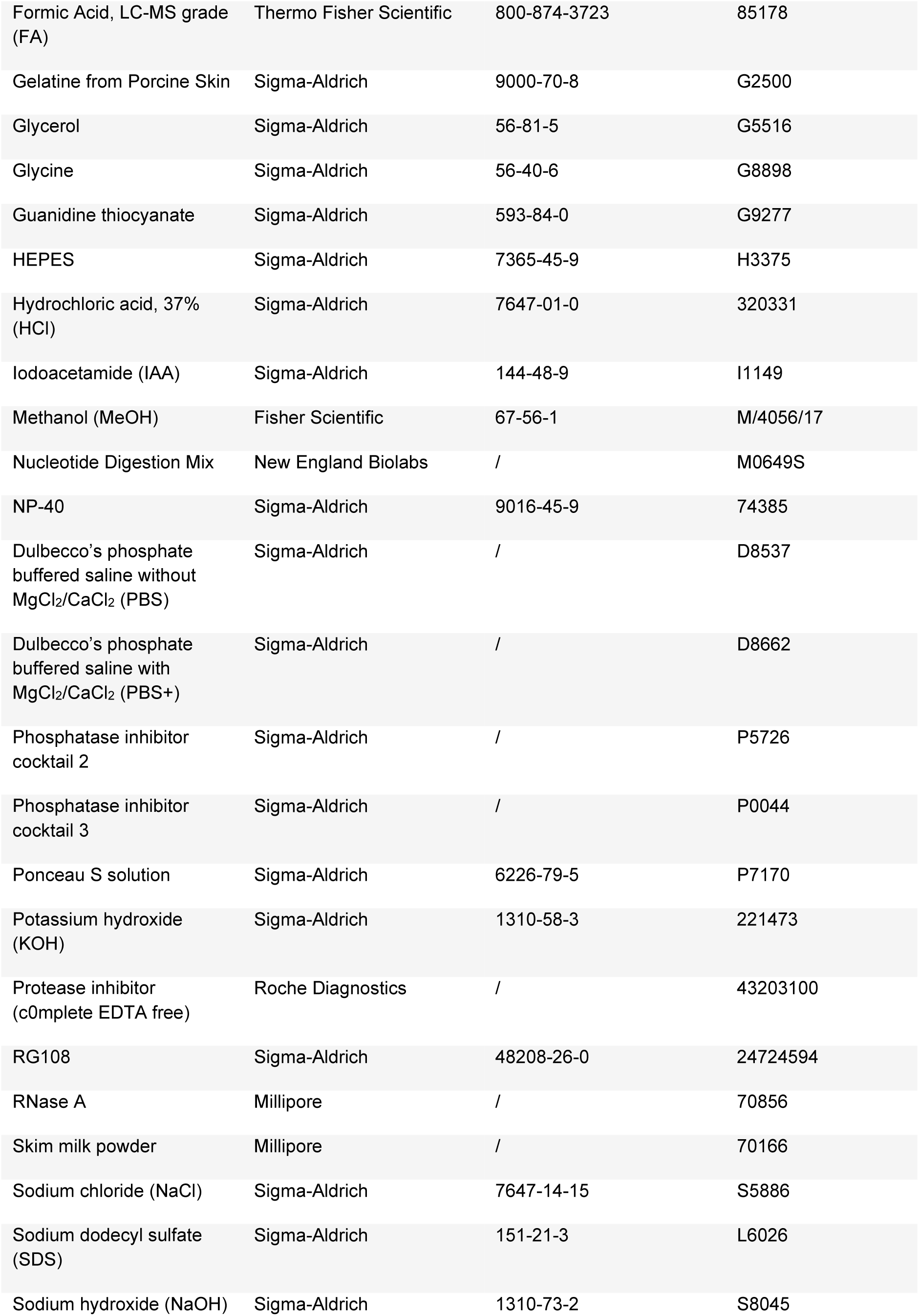
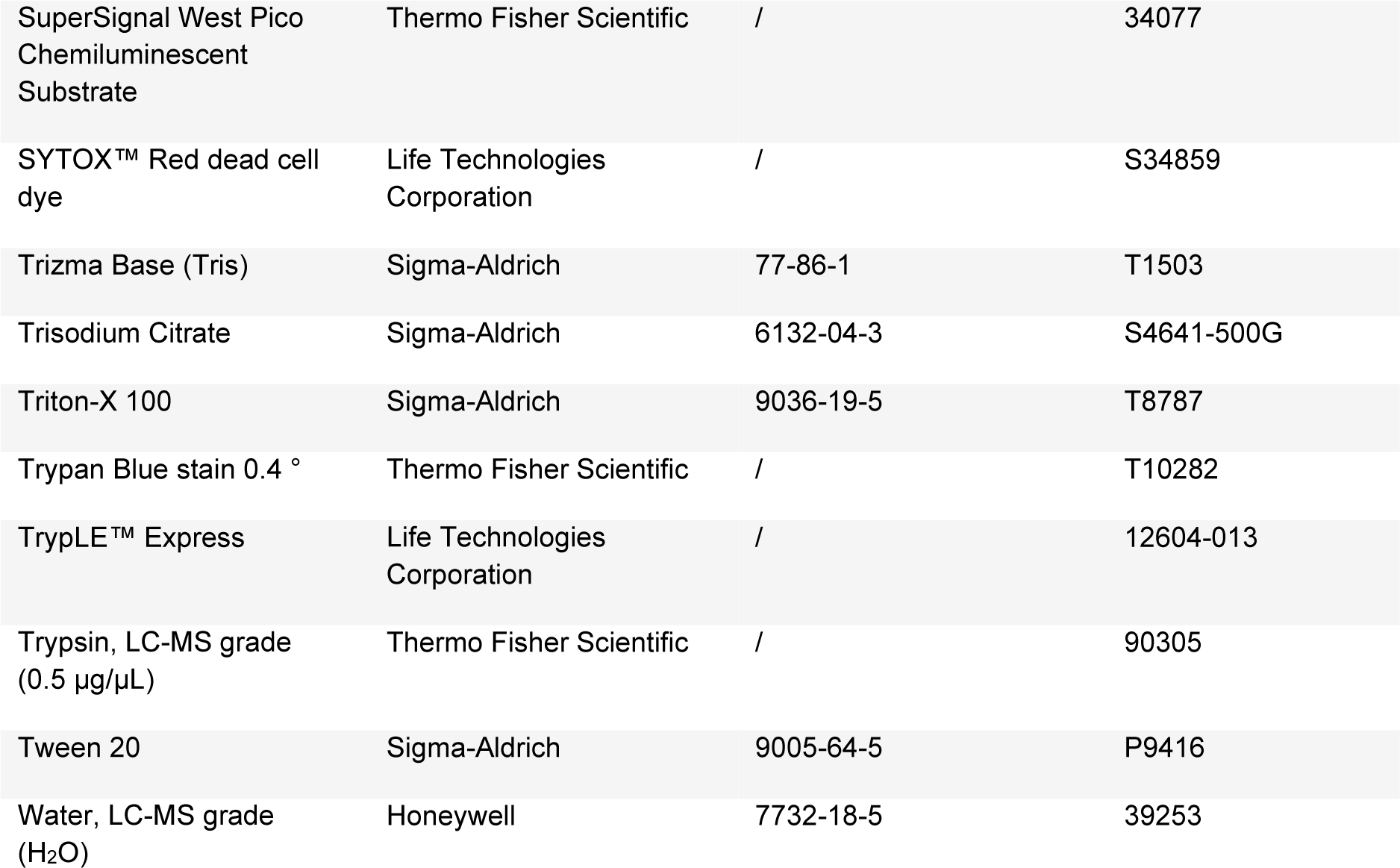
List of chemicals used in this study.

AzadC and AzaC were dissolved in water to a concentration of 10 mM and directly shock-frozen in liquid nitrogen and stored in 10 µL aliquots at −80 °C. Aliquots were only thawed once prior to addition and diluted to a 100 µM solution with water. Integrity of AzadC and AzaC stocks was routinely checked by HPLC, followed by MS of the peak fraction.

RG108 was dissolved in DMSO to a concentration of 100 mM and directly shock-frozen in liquid nitrogen and stored in 2.5 µL aliquots at −80 °C. Aliquots were only thawed once prior to addition and directly diluted to 10 mM using 10% (v/v) EtOH in water.

#### Cell culture media and supplements

For culturing mESCs, the media components as listed in Table 3 were used.

**Table 3:** List of media components for mESC culturing.

| Medium component | Manufacturer | CAS number | Catalogue number |
| --- | --- | --- | --- |
| β-Mercaptoethanol | Sigma-Aldrich | 60-24-2 | F63689 |
| CHIR99021 | Axon Medchem | 252917-06-9 | HY-10182 |
| DMEM – high glucose | Sigma-Aldrich | / | D6429 |
| MEM Non-essential Amino Acid (NEAA) Solution (100x) | Sigma-Aldrich | / | M71145 |
| ESGRO® Recombinant Mouse LIF Protein (mLif) | Sigma-Aldrich | / | ESG1107 |
| L-Alanyl-L-Glutamine | Sigma-Aldrich | 39537-23-0 | G8541 |
| Pansera ES-grade FBS | Pan Biotech | / | P30-2602 |
| PD334581 | Axon Medchem | 391210-10-9 | HY-10254 |
| Penicillin-Streptomycin<br>(Pen-Strep) (100x) | Sigma-Aldrich | / | AP0781 |

Media and supplements were stored according to the available product sheet. Supplements that were shipped in dry form, were solubilized and stored appropriately before use. β-Mercaptoethanol was diluted to a concentration of 50 mM in PBS and stabilized with 35 µM EDTA (pH 8.0). mLif was diluted to 10^6^ U/mL with PBS containing 15% (w/v) BSA, sterile filtered, aliquoted and stored at 4 °C for up to two months. FBS was used without heat inactivation.

The mESC basic medium consisted of DMEM, 10% (v/v) FBS, 2 mM L-alanyl-L-glutamine, 0.1 mM β-mercaptoethanol, 1x MEM-NEAA, 1x Pen-Strep. After preparation, the basic medium was sterile filtered using a bottle top filter unit (Ø 0.2 µm). To prepare Lif medium from basic medium, mLif was added to a final concentration of 10^3^ U/mL. To prepare 2iL medium from basic medium, mLif was added to a final concentration of 10^3^ U/mL, CHIR99021 and PD334581 were added to a final concentration of 3 µM. Basic and Lif media were stored for up to two weeks at 4 °C, 2iL medium was stored for up to ten days at 4 °C.

#### Antibodies

All antibodies used in this study are listed in Table 4.

**Table 4:** List of antibodies used in this study. mAB = monoclonal antibody; pAB = polyclonal antibody.

| Antibody | Manufacturer | Catalogue number | Application |
| --- | --- | --- | --- |
| anti-phospho Histone H2A.X (Ser139), clone JBW301 (mouse mAB) | Millipore | 05-636 | Western blotting (1:1000), Immunofluorescence (1:250) |
| Anti-Histone H3, clone D1H2 (rabbit mAB) | Cell Signaling | 4970S | Western blotting (1:1500) |
| Anti-PARP1 (rabbit pAB) | Proteintech | 13371-1-AP | Western blotting (1:1000) |
| Alexa Fluor® 488-conjugated AffiniPure Rat Anti-Mouse IgG (H+L) | Jackson ImmunoResearch | 415-545-166 | Immunofluorescence (1:400) |
| HRP-conjugated AffiniPure Goat Anti-Mouse IgG (H+L) | Proteintech | SA00001-1 | Western blotting (1:7500 – 1:10000) |
| HRP-conjugated AffiniPure Goat Anti-Rabbit IgG (H+L) | Proteintech | SA00001-2 | Western blotting (1:7500 – 1:10000) |

### Biological Resources

wt J1 mESCs were described in *Li et al., 1992* (33) and originally provided by the Jaenisch lab (Whitehead Institute, USA). DNMT-TKO J1 mESCs were described in *Tsumara et al., 2006* (28) and originally provided by the Okano lab (RIKEN, Japan). Trdmt1-KO J1 mESCs were described in *Okano et al., 1998* (34) and originally provided by the Jaenisch lab.

### Statistical Analysis

For the proteomics experiments, details about the statistical analysis, including sample size, data exclusion and significance thresholds, are given in the Proteomics – Materials and Methods section, including the respective analysis software. For all other experiments, Graph Pad Prism (v 9.4.0)was used for statistical analysis and details are given in the Supplementary

Data File 1. No statistical methods were used to pre-determine sample size. Sample sizes were chosen based on cost, experience and commonly used sample sizes for *in vitro* experiments (n ≥ 3), which provided in this study low inter-sample variety between samples of the same group. The number of samples for each experiment is either described in the Materials and Methods section or directly evident from the respective figure and figure legend.

### Data Availability

Original (unprocessed) and metadata are deposited as described in the Data Availability Section.

### Data Base References

We used the reactome knowledge data base (https://reactome.org, access March 2023 – August 2023; (35)) to assign DNA repair-relevant proteins in our proteomics data sets. To receive optimal coverage, we mapped the mouse proteins to the respective human proteins via the gene name and did the subsequent analysis on the human pathways.

The UniProt database (https://uniprot.org) was used to download the FASTA file of *mus musculus*.

GOrilla (https://cbl-gorilla.cs.technion.ac.il/; access February 2023, (36)) was used for Gene Ontology (GO) term analysis.

### mESCs Handling

#### Culture conditions and passaging

mESCs were cultivated at 37 °C in water saturated, CO_2_-enriched (5%) atmosphere on gelatine-coated plates. For gelatine coating, 0.2% (w/v) gelatine in water was prepared, heat sterilized, brought to RT and filtered. Afterwards, culture dishes were coated for 10 to 60 min at 37 °C, coating solution was aspirated and mESCs were directly plated in the appropriate amount of medium. For mESC maintenance, 2iL medium was used as a standard medium. mESCs were routinely passaged every 2 – 3 d in a ratio of 1:4 to 1:8 when reaching a confluency of 60 – 75%. To detach the mESCs, medium was aspirated, cells were washed with PBS and TrypLE (150 µL/6well) was added for 4 – 5 min at 37 °C before trypsination was stopped with medium. Then, mESCs were resuspended to a single cell solution, the required amount of cells was centrifuged at 300 × g for 3 min at RT and afterwards replated in new medium. When reaching passage #25 after thawing, cells were discarded. mESCs were checked once during cultivation for Mycoplasma contamination using a PCR-based mycoplasma detection kit (Jena Bioscience #PP-401L) as indicated by the manufacturer.

#### Priming

For priming, the anticipated portion of mESCs were cultured after passaging in Lif medium instead of 2iL. After 48 h, mESCs on Lif medium were passaged again in new Lif medium. If not indicated otherwise, cells were primed for 96 h in total before analysis.

#### Treatment

Experiments were only started if cell and colony morphology indicated high cellular fitness and cell viability was > 90% as indicated by Trypan Blue staining. In all experiments, the “Untreated Ctrl.” refers to mESCs of the same genotype for a respective experiment, which was not treated with any compound the same way otherwise. For each experiment, untreated and treated samples were seeded from the same mESC batch and then handled in parallel.

The number of seeded cells depended on plate size: ca. 115,000 cells for 12 well, ca. 300,000 cells for a 6well, ca. 700,000 cells for a p60, ca. 1,900,000 cells for a p100 and ca. 4,500,000 cells for a p100. AzadC and AzaC was added at the indicated concentrations from the diluted 100 µM stock solution directly into the medium after seeding. Unless stated otherwise, medium was not changed anymore until cell harvest 48 h after start of the treatment. For treatment under naïve conditions, 2iL medium was used (2iL, 48 h treatment). For treatment under primed conditions, mESCs were primed for 48 h before treatment and the compounds were added to Lif medium after the second passaging in Lif medium (in total 96 h primed in Lif, treatment in the last 48 h). For RG108 experiments, mESCs were maintained for 4 weeks in 2iL medium containing 50 µM of RG108. For priming, mESCs were seeded for 48 h in Lif medium containing 50 µM of RG108 before they were finally seeded into Lif medium containing 200 µM of RG108 for additional 48 h.

#### Proliferation assay

For the proliferation assay, two wells were seeded per biologically independent sample type. After 24 h, the first well per sample was harvested and cells were counted and after 48 h, the second well per sample was harvested and the cells were counted. Counting was done using a Countess 3^TM^ automated cell counter (Invitrogen).

### Microscopy

#### Brightfield microscopy

For brightfield microscopy images, cells were imaged directly in the medium at the end of treatment using an EVOS^TM^ M5000 imaging system in transmission mode and 10x or 20x magnification. Afterwards, brightness and contrast were automatically adjusted using Adobe Photoshop 2023 for optimal visualization of the cells. Pictures were taken from representative regions.

#### Fluorescence confocal microscopy

All steps were performed in a humidity chamber and at RT if not otherwise specified. 30,000 cells per well were seeded in an ibidi µ-slide 8 well (ibidi #80826) and treated as indicated. After 48 h treatment, cells were washed with PBS+ and fixed for 10 min using 4% formaldehyde solution. After three times washing with PBS+, the cells were permeabilized and blocked for 30 min using 0.3% (v/v) Triton X-100 and 5% (v/v) Chemiblocker. The primary anti-gH2A.X antibody was diluted in PBS+, containing 5% (v/v) CB and 0.3% (v/v) Triton X-100 and applied over night at 4 °C. After incubation, mESCs were washed three times with PBS+ containing 2% (v/v) CB. For secondary detection, the fluorescent labeled Alexa488 anti-mouse antibody was diluted in PBS+, containing 3% (v/v) CB and applied for 1 h in the dark, followed by three times washing with PBS+. Cell nuclei were stained with Hoechst 33342 (5 µg/mL), which was applied for 15 min in the dark, followed by one washing step with PBS+. After mounting, the samples were analyzed using a Leica SP8 confocal laser scanning microscope with the associated LAS X software (Leica, Wetzlar). Regions for imaging were chosen based on Hoechst signal. Brightness and contrast were adjusted for the control using ImageJ (version 1.54a) and afterwards the settings were applied to all other images.

### Triple-Quadrupole Mass Spectrometry (QQQ-MS) for Nucleoside Quantification

For QQQ-MS experiments, mESCs were seeded in a p60 and treated with the indicated concentrations. For the AzadC and AzaC incorporation experiments, cells were either treated for 24 h (treatment a), 48 h without medium change (treatment b), 48 h with medium change after 24 h without second compound addition (treatment c) or 48 h with medium change after 24 h and second compound addition at the same concentration (treatment d). For all other QQQ-MS experiments, mESCs were treated for 48 h without medium change (treatment b). After treatment was finished, medium was aspirated, mESCs were washed with PBS and harvested. After harvest, mESCs were washed once with PBS and afterwards directly lysed in 800 µL of GTC buffer (3.5 M guanidine thiocyanate, 25 mM trisodium dihydrate, 14.3 mM β-mercaptoethanol, pH 6.9) and either processed directly or shock-frozen in liquid N_2_ and stored at −80 °C. For thawing, lysed samples were quickly warmed to RT and gDNA isolation was performed.

#### gDNA and RNA isolation

After cell lysis, gDNA and RNA isolation was performed according to *Traube et al., 2019* (37) with minor modifications. Butylated hydroxytoluene and deferoxamine were not added to the washing buffers and mESCs were only lysed by adding the chaotropic GTC buffer, but the bead mill step described in the previously published protocol was skipped. For AzadC and AzaC incorporation experiments, isolated gDNA or RNA was directly subjected to nucleoside digest and QQQ-MS measurements. For mdC quantification, the isolated gDNA could be stored at −80 °C before nucleotide digestion.

#### Nucleotide digestion

Nucleotide digestion was performed in technical duplicates per biologically independent sample. Two digestion controls were added for each digestion as described in *Traube et al., 2019* (37). Per sample, 0.5 µg of gDNA or RNA was digested in a total volume of 30 µL using 0.5 µL of enzyme and 3 µL of 10x buffer from the Nucleotide Digestion Mix for 1 h at 37 °C. Afterwards, 20 µL of water were added to reach a final volume of 50 µL. For AzadC and AzaC incorporation experiments, samples were filtered immediately as described in *Traube et al., 2019* (37). After filtration, the samples were directly subjected to QQQ-MS. For mdC quantification, the nucleoside mixture could be stored at −20 °C before filtering and analyzing.

#### QQQ-MS data acquisition

For QQQ-MS, an Agilent 1290 Infinity equipped with a variable wavelength detector (VWD) combined with an Agilent Technologies G6490 Triple Quad LC/MS system with electrospray ionization (ESI-MS, Agilent Jetstream) was used. All solvents were LC-MS grade. The operating parameters were: positive-ion mode, cell accelerator voltage of 5 V, N_2_ gas temperature of 120 °C and N_2_ gas flow of 11 L/min, sheath gas (N_2_) temperature of 280 °C with a flow of 11 L/min, capillary voltage of 3000 V, nozzle voltage of 0 V, nebulizer at 60 psi, high-pressure RF at 100 V and low-pressure RF at 60 V. The instrument was operated in dynamic MRM mode (Supplementary Data File 2, Table S1 – Table S3). For separation, an Poroshell 120 SB-C8 column (2.7 μm, 2.1 × 150 mm; Agilent Technologies, #683775-906) was used. Running conditions were 35 °C and a flow rate of 0.35 mL/min for all experiments. Specifications for AzadC incorporation experiments were (Supplementary Data File, Table S1): binary mobile phase of 5 mM NH_4_OAc aqueous buffer A (pH 5.3) and an organic buffer B of 0.0075% FA in MeCN. The gradient started at 100% solvent A for 1.5 min, followed by an increase of solvent B to 20% over 7 min (1.5 min – 8.5 min) and further to 80% B within the following minute (8.5 min – 9.5 min). 80% B was maintained for 2.5 min (9.5 min – 12 min) before returning to 100% solvent A in 0.5 min and a 2.2 min re-equilibration period. Specification of AzaC incorporations were (Supplementary Data File, Table S2): 0.0075% FA in aqueous buffer A and an organic buffer B of 0.0075% FA in MeCN. The gradient started at 100% solvent A for 1.2 min, followed by an increase of solvent B to 5% over 5 min (1.2 min – 6.2 min) and further to 80% B within the following 1.3 min (6.2 min – 7.5 min). 80% B was maintained for 2 min (7.5 min – 9.5 min) before returning to 100% solvent A in 0.5 min and a 2.5 min re-equilibration period. Specification for mdC quantification were (Supplementary Data File, Table S3): binary mobile phase of 0.0075% FA in aqueous buffer A and an organic buffer B of 0.0075% FA in MeCN. The gradient started at 100 % solvent A, followed by an increase of solvent B to 3.5% over 4 min (0 min – 4 min) and from 4 min to 7 min solvent B was further increased to 5%. From 7.0 min to 8.0 min, solvent B was increased to 80 % and maintained at 80 % for 2.5 min before returning to 100 % solvent A in 1.5 min and a 2.2 min re-equilibration period. Of each sample, 10 μL were co-injected with 1 μL of stable isotope labeled internal standard (ISTD). The ISTD mix consisted for all measurements of 200 µM theophylline to have an ISTD mix-UV control, and of 0.5 µM of each isotope standard (^15^N_5_-^13^C_10_-dA, ^13^C_9_-dC, ^15^N_5_-^13^C_10_-dG, ^15^N_2_-^13^C_5_-dT, D_3_-m^5^dC, ^15^N_2_-D_2_-hm^5^dC, ^15^N_2_-f^5^dC, ^15^N_2_-ca^5^dC and ^15^N_5_-8oxodG). Calibration curves for canonical nucleosides (dA, dC, dG and dT) spanned 0.1 pmol – 200 pmol and for the modified nucleosides (mdC, hmdC, fdC, cadC and 8oxodG) 0.004 pmol – 5 pmol.

#### Analysis

The sample data were analyzed by the quantitative and qualitative MassHunter Software from Agilent (v B07.01). As there was no internal standard available for exact quantification of AzadC and AzaC, we calculated the area under the curve ratio for AzadC/dG or for RNA measurements, the AzaC/G ratio to obtain the relative incorporation levels normalized to the dG and G content. For mdC quantification, we followed the procedure as described in *Traube et al. 2019* (37), except that also the amount of canonical nucleosides were calculated by MS and not via the UV-trace. Samples, where the sum of C-modifications (dC + mdC + hmdC) deviated by more than 15% from dG were discarded as since they did not pass the quality threshold. As we measured each sample in technical duplicates, we calculated the mean of the technical replicates to obtain the mean for each biologically independent sample.

### Flow Cytometry-based Apoptosis Assay

For flow cytometry, mESCs were seeded in a 12well and treated as indicated. Before harvest, the medium, which includes dead floating cells, was not aspirated but collected as well and the remaining attached cells were harvest and combined with the cells from the medium. Afterwards, mESCs were washed twice with PBS and subsequently counted. 1.5 × 10^5^ cells per sample were transferred into a new tube. Apoptosis and necrosis were determined by using the FITC Annexin V Apoptosis Detection Kit and SYTOX^TM^ Red Dead Cell Stain. To this end, cells were resuspended in 150 µL of Annexin V binding buffer supplemented with 0.75 µL of FITC-conjugated Annexin V and 0.15 µL of SYTOX^TM^ Red Dead Cell Stain, gently vortexed and incubated at RT for 15 min in the dark. Afterwards, samples were put on ice and the cell suspension was filtered through a 35 µm strainer prior to measurement. For the analysis, BD FACSCanto^TM^ (recording of 10,000 events per sample) and FlowJo Single Cell Analysis Software (v10.8.0) were used. Gates (FSC (A) – SSC (A) to remove cell debris, FSC (A) – FSC (H) to gate for single cells and last FITC/APC to distinguish between live, dead and early apoptotic cells) were set once for the control sample and then applied to all other samples.

### Immunoblot analysis

#### Nuclear extract preparation

For the preparation of nuclear extracts, which were used for western blotting to enrich nuclear-specific proteins, mESCs were seeded in a p100 and treated for 48 h. After treatment, the mESCs were harvested and nuclear extracts were prepared as previously described by *Dignam et al., 1983* (38) with the modification that every buffer was supplemented with phosphatase inhibitor cocktail 2 and phosphatase inhibitor cocktail 3, 1:100 each. Furthermore, c0mplete protease inhibitor was used to inhibit any protease activities. Afterwards, the protein concentration was determined using a Bradford assay as described by the manufacturer. SDS loading buffer (final concentration 50 mM Tris-HCl (pH 6.8), 100 mM DTT, 2% (w/v) SDS, 10% (v/v) glycerol, 0.25% (w/v) bromophenol blue) was added. The samples were vortexed, incubated for 5 min at 92 °C and afterwards stored at −20 °C. Before loading the samples on a polyacrylamide gel, the samples were heated for additional 2 min at 92 °C and vortexed thoroughly.

#### Western blotting (Immunoblot)

Per sample, 15 µg of nuclear extract in SDS loading buffer were loaded on a 4-15% precast polyacrylamide gel (Bio-Rad #4561083EDU) and Color-coded Prestained Protein Marker, Broad Range (10-250 kDa) (New England Biolabs #P7719S) was used as a protein standard. The gel was run at constant 150 V for 60 min in SDS running buffer (25 mM Tris, 192 mM glycine, 0.1% (w/v) SDS). For blotting, we used a PVDF blotting membrane (GE Healthcare Amersham Hybond P0.45 PVDG membrane #10600023) and pre-cooled Towbin blotting buffer (25 mM Tris, 192 mM glycine, 20% (v/v) MeOH, 0.038% (w/v) SDS). The membrane was activated for 1 min in methanol, washed with water and equilibrated for additional 2 min in Towbin blotting buffer; the Whatman gel blotting papers (Sigma-Aldrich #WHA10426981) were equilibrated for 15 min in Towbin buffer and the precast gel was equilibrated for 5 min in Towbin buffer after the run. Western blotting (tank (wet) electro transfer) was performed at 4 °C for 9 h at constant 35 V. After blotting, the PVDF membrane was blocked for 1 h at RT and constant shaking using 5% (w/v) skim milk powder in TBS-T (20 mM Tris-HCl (pH 7.5), 150 mM NaCl, 0.1% (v/v) Tween-20). The primary antibodies were diluted in 5 mL of 5% (w/v) skim milk powder in TBS-T. The blocking suspension was discarded, and the diluted primary antibodies were added for 12 - 16 h at 4 °C and shaking. After incubation, the primary antibodies were discarded, and the membrane was washed three times 10 min with TBS-T. HRP-conjugated secondary antibodies were diluted in 5% (w/v) milk powder in TBS-T and added for 1 h at room temperature under shaking. Afterwards, the membrane was washed two times with TBS-T and one time with TBS (TBS-T without Tween-20) before SuperSignal West Pico Chemiluminescent Substrate was used for imaging. Western blots were imaged using Amersham Imager 680 (auto exposure mode).

For imaging the same blot multiple times using different antibodies, the membrane was directly stripped after imaging. To this end, the membrane was put in TBS-T and the buffer was heated in a microwave until boiling. Afterwards, the buffer was discarded and the procedure was repeated in total three times. After stripping, the membrane was blocked again using 5% (w/v) milk powder in TBS-T and the protocol followed the above described procedure.

At the end, the membrane was stained with Ponceau S to visualize the total protein load.

### Liquid-chromatography tandem mass spectrometry (LC-MS/MS)) – Proteomics

For proteomics experiments, mESCs were seeded in a p150 and treated for 48 h. For each sample type (specific combination of genotype, culturing conditions, treatment), four biologically independent replicates were initially generated for chromatin enrichment and at least three biologically independent replicates were generated for whole proteome analysis. After treatment, mESCs were harvested and washed with PBS. 10% of the cells were used for whole proteome isolation and the rest for chromatin enrichment to generate whole proteome and chromatin enriched proteome samples from the same batch of cells.

#### Whole proteome isolation

Cells were lysed in 200 µL of total lysis buffer (20 mM HEPES, 1% (v/v) NP-40 and 0.2% (w/v) SDS for 30 min on ice and afterwards centrifuged at 21,000 × g at 4 °C for 10 min. Supernatant containing the proteins was transferred to a new tube.

#### Chromatin enrichment

Chromatin extraction was performed according to *Gillotin, 2018* (39). The cell pellet was lysed in 150 µL (ca. 5x pellet size) of E1 lysis buffer containing 50 mM HEPES-KOH, 140 mM NaCl, 1 mM EDTA, 10% (v/v) glycerol, 0.5% (v/v) NP-40, 0.25% (v/v) Triton-X 100, 1 mM DTT and c0mplete protease inhibitor. The cell extract was then centrifuged at 1100 × g, 4 °C for 2 min and the supernatant (cytoplasmic fraction) was transferred into a new tube. The pellet was resuspended again in E1 lysis buffer and incubated for 10 min on ice and then centrifuged again at 1100 × g, 4 °C for 2 min. Supernatant was discarded. Next, the pellet was washed three times in 50 µL ice-cold E2 buffer containing 1 M Tris-HCl (pH 8.0), 200 mM NaCl, 1 mM EDTA, 0.5 mM EGTA and c0mplete protease inhibitor, by resuspending the pellet fully and centrifuging at 1100 g, 4 °C for 2 min. The supernatants (nuclear fraction) were transferred and pooled in a new tube. In the last of the three washing steps, the sample was incubated on ice for 10 min before centrifugation. After centrifugation, the remaining pellet was resuspended in E3 buffer containing 1 M Tris-HCl (pH 7.5), 20 mM NaCl, 1 mM MgCl2, 0.1% (v/v) Benzonase and c0mplete protease inhibitor. The pellets were resuspended in this buffer, sonicated for 10 cycles with 30 ‘’ on and 30 ‘’ off at maximum power at 4 °C using a Bioruptor Pico sonication device (Diagenode). Afterwards, the samples were centrifuged at 16000 × g, 4 °C for 10 min. The supernatant containing the chromatin bound proteins was transferred into a new tube for further analysis.

Protein concentrations were determined by bicinchoninic acid assay (BCA) using the BCA protein assay kit according to the manufacturer’s protocol. Every sample was measured in technical duplicates at 562 nm on a multimode microplate reader (Tecan) and the mean of the two technical replicates was calculated for each sample. The total protein concentration was calculated using a calibration curve prepared with BSA that was re-done for every measurement. Because of poor quality and low protein amount, one sample of the whole proteome fraction for each sample (2iL and Lif) could not be analyzed and had to be discarded, leaving three biologically independent replicates per sample type for whole proteome measurement.

#### SP3 protocol

20 µg of protein were used for each sample. The protein sample was added to pre-washed Dynabeads™ Protein G (bead/protein ratio 10:1, ca. 7 µL of resuspended beads for 200 µg) and filled up with total lysis buffer to a working volume of 50 µL. The samples were incubated on the beads while shaking at 1000 rpm for 1 min at RT. 120 µL of EtOH were added and then the samples were incubated again for 5 min, shaking at 1000 rpm. The beads were trapped on a magnet for 2 min and the supernatant was discarded. The bead-bound proteins were then washed three times with 100 µL 80% (v/v) ethanol, incubated for 1 min while shaking at 850 rpm for each step and then supernatant was discarded on a magnet. The beads were resuspended in 100 µL of 100 mM ABC buffer, then 10 mM DTT and 20 mM IAA were added from a 1 M stock solution, respectively and the samples were incubated for 5 min at 95 °C, shaking at 850 rpm. After the samples were cooled down to RT, Trypsin protease (LC-MS grade) was added to the sample at a ratio of 1:50 (0.4 µg of Trypsin for 20 µg of protein) and incubated over night at 37 °C, shaking at 850 rpm. The peptides were then carefully transferred into a fresh tube and the beads were washed twice with 50 µL 0.1% (v/v) FA. The three fractions were pooled and again incubated on a magnet and transferred into a fresh tube to remove all beads from the sample. Then, the peptide samples could be stored at −80 °C until further analysis.

#### MS acquisition and analysis

MS analysis was performed as described in *Makarov et al., 2022* (40) on an Orbitrap Eclipse Tribrid Mass Spectrometer (Thermo Fisher Scientific) coupled to an UltiMate 3000 Nano-HPLC (Thermo Fisher Scientific) via an EASY-Spray source (Thermo Fisher Scientific) and FAIMS interface (Thermo Fisher Scientific) using data-independent acquisition (DIA) mode. LC-MS grade solvents were used. Per sample, 1 µg of peptides were first loaded on an Acclaim PepMap 100 μ-precolumn cartridge (5 μm, 100 A, 300 μm ID x 5 mm, Thermo Fisher Scientific) and were then separated at 40 °C on a PicoTip emitter (noncoated, 15 cm, 75 μm ID, 8 μm tip, New Objective) that was *in-house* packed with Reprosil-Pur 120C18-AQ material (1.9 μm, 150 Å, Dr. A. Maisch GmbH). A gradient over 60 min and 0.1% (v/v) FA in LC-MS grade water as buffer A and 0.1% (v/v) FA in MeCN as buffer B were used with a flow rate of 0.3 µL/min and 0 – 5 min 4% B, then from 5 min – 6 min to 7% B, followed by 6 min – 36 min to 24.8% B, 36 min – 41 min 35.2% B. From 41 min – 41.1 min B was increased to 80% until 46 min, when column was re-equilibrated at 4% B until 55 min. The DIA duty cycle consisted of one MS1 scan followed by 30 MS2 scans with an isolation window of the 4 m/z range, overlapping with an adjacent window at the 2 m/z range. MS1 scan was conducted with Orbitrap at 60,000 resolution power and a scan range of 200 - 1,800 m/z with an adjusted RF lens at 30%. MS2 scans were conducted with Orbitrap at 30,000 resolution power, RF lens was set to 30%. The precursor mass window was restricted to a 500 – 740 m/z range. HCD fragmentation was enabled as activation type with a fixed collision energy of 35%. FAIMS was performed with one CV at −45 V for both MS1 and MS2 scans during the duty cycle.

#### Analysis of MS spectra and pathway analysis

MS raw files were processed in DIA-NN (v. 1.8.1) (41) to determine the protein identity and quantity in each sample. FASTA digest for library-free search/library generation was activated, and a FASTA spectral library generated for *mus musculus* from UniProt (uniport.org) was used. As protease, Trypsin/P was chosen and the missed cleavages being allowed were set to 2 and the minimal peptide length was set to 6. MBR (match between runs) mode was enabled. As quantification strategy Robust LC (high precision) was used and threads were set to 7. Otherwise default settings were used. From the DIA-NN output, the protein groups file was uploaded to and subsequently analysed using MaxQuant Perseus (v. 1.6.15) (42). First, samples where the number of identified unique proteins (1% FDR) was <500, were discarded to avoid analysis bias by many missing values that would have to be imputed in the further analysis steps. Next, samples were group according to genotype, culturing condition and treatment (samples that are biologically independent but of the same type). Proteins had to be present in at least >50% of the samples within one sample group or were otherwise completely discarded from the analysis to avoid bias by a large number of missing values. For the remaining proteins, LFQ intensities were log2 transformed and missing values were replaced from normal distribution (separately for each column, default settings). The wt and the DNMT-TKO samples were separately analysed using the Volcano plot function in Perseus (t-test, both sided, 250 randomizations, FDR 0.05) and treated samples were analysed against the respective controls. We applied a stricter significance threshold regarding the log_2_FC for the chromatin-enriched proteins with a significance threshold of −log(p-value) > 1.3 and |log_2_FC| > 1 (|fold change| > 2) compared to the whole proteome where a significance threshold of −log(p-value) > 1.3 and |log_2_FC| > 0.58496 (|fold change| > 1.5) was applied, because more preparation steps were required to isolate the chromatin-bound proteins. This can increase inter-sample variation independent from biological reasons. Furthermore, we wanted to ensure to take only proteins into account for the downstream analysis that showed a very strong chromatin-recruitment as a response to AzadC treatment.

Next, DNA repair proteins were assigned according to reactome (35) projecting the mouse proteins to the human equivalents. For identifying enriched pathways within the proteomics data sets, the pathfindR tool was used as described (43). Pathways were considered as significantly enriched when the criteria fold enrichment ≥ 2, highest p-value < 0.05 were both fulfilled. GO analysis was performed using Gorilla (36) to analyse the chromatin fraction against the whole proteome of untreated wt cells with the settings specified in Supplementary Data File 3.

## RESULTS

### mESCs as a model system to study the mode of action of AzadC and AzaC

Before starting to investigate the effects of AzadC and AzaC using our anticipated model system and before drawing conclusions from it, we had to ensure that the wt and the DNMT-TKO cells provided similar cellular features, e.g. similar morphological changes upon priming, despite the absence of all three DNMT enzymes. When cultured in 2iL, both, wt and DNMT-TKO cells, formed densely packed colonies with distinct borders. Within the colonies, individual cells could not be spotted (Figure 1B, naïve). Upon priming, the colonies dissected and individual cells with distinct morphology were increasingly observed for both genotypes (Figure 1B, primed). Moreover, a comparable number of proteins showed significant expression level changes from the naïve to the primed state for both genotypes (Figure 1C). Although the overall protein expression levels of both genotypes were different in the two genotypes (Figure 1D, Figure S1B), the expression changes of the common differentially expressed proteins between the naïve and the primed state showed a very high correlation (Pearson r = 0.8744, Figure 1D). Altogether, these results confirmed that the wt and the DNMT-TKO mESCs undergo fundamental and comparable cellular changes from the naïve to the 96 h primed state and were therefore a suitable model system to directly compare the effects of AzadC and AzaC in the presence and absence of DNMT enzymes. Furthermore, the wt also provides a suitable system to study the effects of AzadC and AzaC under low and high DNMT activity. Upon 96 h priming, Dnmt1, Dnmt3a and Dnmt3b were all significantly higher expressed on the protein level compared to the naïve state (Figure 1E), which consequently resulted in higher DNMT activity as indicated by increasing levels of 5mdC (Figure 1F). Since new 5mdC patterns are established during priming, wt mESCs allow to monitor closely the effects of AzadC and AzaC when DNMT activity is highly dynamic.

Based on these observations, we concluded that the wt and the DNMT-TKO represent an ideal model system to distinguish in a combinatorial approach the DNMT-based effects of AzadC and AzaC from DNMT-independent effects (Figure 1G, Table 1). It is expected that DNA-DNMT crosslinking, which can only occur in the wt, has a major impact on AzadC- or AzaC-treated cells because it results in DNA hypomethylation (MoA 1) and deleterious DNA double strand breaks (MoA 2). Using our mESC model system, we were able to additionally distinguish between the DNMT-DNA crosslinking effects under low DNMT activity (MoA 1a/MoA 2a) and under high DNMT activity (MoA 1b/MoA 2b). DNA lesions that occur from base pairing mismatches (MoA 3) or creation of abasic sites from hydrolysis (MoA 4) after exposure to AzadC or AzaC is expected to have minor effects but should be a burden for wt and DNMT-TKO cells. Furthermore, the DNMT-TKO allows to study the RNA-based effect of AzaC in more detail without interferences from DNA-DNMT crosslinking. As the substantial changes on the proteome level during priming require drastic changes of RNA-transcription and -translation, the RNA-dependent effect of AzaC should be equally observable in the wt and the DNMT-TKO.

### The non-canonical nucleobase 5-aza-cytosine substantially contributes to the anti-proliferative and cytotoxic effect of AzadC when DNMT-activity is low

To investigate how much the non-canonical nucleobase 5-aza-cytosine contributes to the toxicity of AzadC by base mismatch (MoA 3) or formation of abasic sites by spontaneous hydrolysis of the base (MoA 4), independent from DNA hypomethylation (MoA 1) and DNA damage by DNMT-crosslinking (MoA 2), we first compared the effects of AzadC on wt and DNMT-TKO cells under naïve conditions (2iL) when DNMT-expression and -activity are low in the wt. First, we confirmed by our previously reported QQQ-MS method for exact quantification of nucleosides (37) that AzadC had the anticipated hypomethylating effect in the wt even under low DNMT-activity (Figure 2A). We detected a significant decrease in the amount of mdC after 48 h treatment with 1.25 µM or 2.5 µM of AzadC with no difference between the concentrations, indicating that the hypomethylating effect was already at the maximum at the concentrations applied. Since the mdC levels of the DNMT-TKO were under the limit of detection (LOD) even without treatment (Figure S1D), a hypomethylating effect could not be observed upon AzadC treatment. Even though DNA-demethylation was at the maximum at 1.25 µM AzadC in the wt, inspection of the phenotypic changes of the cells by brightfield microscopy revealed that the anti-proliferative and cytotoxic effect of AzadC was strictly concentration-dependent with only mild effects at 1.25 µM of AzadC in the wt (Figure 2B). This result was in line with previous reports that AzadC-induced DNA hypomethylation cannot be used as a prognostic marker for induction of cell death by AzadC (44). Interestingly, the DNMT-TKO showed a very similar cell viability pattern, albeit starting at higher concentrations (Figure 2B). Next, we quantified the proliferation rate of the wt and the DNMT-TKO when the cells were exposed to 2.5 µM of AzadC and compared it to untreated cells (Figure 2C). The untreated wt and DNMT-TKO cells showed an identical proliferation rate with a doubling time of about 17 h, whereas treatment with 2.5 µM of AzadC resulted in both genotypes in a significantly slower proliferation rate of 26 h doubling time, with again no difference between the wt and the DNMT-TKO cells. The similarity of phenotypic (Figure 2B) and proliferative (Figure 2C) changes in the wt and DNMT-TKO after 48 h treatment with AzadC was highly unexpected because AzadC cannot induce deleterious DNMT-DNA crosslinks in the DNMT-TKO. Consequently one would expect a substantially higher impact of AzadC on the wt. There were two possible options to explain those results, either AzadC is metabolized differently in the wt and the DNMT-TKO, with higher uptake and genomic incorporation in the DNMT-TKO, or the non-canonical 5-azacytosine nucleobase contributes substantially to the anti-proliferative and cytotoxic effects of AzadC when overall DNMT-activity is low. To test which option explained our results, we checked AzadC uptake and genomic incorporation in the wt and the DNMT-TKO by treating both genotypes with 2.5 µM of AzadC for 24 h and 48 h and quantifying the global, sequence context-independent, genomic incorporation levels of AzadC by QQQ-MS (Figure 2D). For the 24 h timepoint, we added 2.5 µM of AzadC once at the start (0 h) and harvested the cells after 24 h (treatment a). For the 48 h timepoint, we treated the cells for 48 h before harvest, but tested three different treatment regimens – addition of 2.5 µM of AzadC at 0 h and neither medium change nor second compound addition (treatment b), addition of 2.5 µM of AzadC at 0 h and medium change after 24 h but no second compound addition (treatment c), and addition of 2.5 µM of AzadC at 0 h and medium change after 24 h with second compound addition of 2.5 µM of AzadC (treatment d). The untreated control served as a background control and as expected, an AzadC signal was not detectable neither in the wt nor in the DNMT-TKO (Figure S2A). For all timepoints and treatment regimens tested, we did not observe any difference between the wt and the DNMT-TKO (Figure 2D), indicating that there was no difference between the two genotypes regarding AzadC uptake and metabolization. However, we detected substantial differences between the different treatment regimens. After 24 h, the amount of genomically incorporated AzadC, normalized to the amount of dG, was about two times higher compared to the 48 h timepoint when no additional compound was added. In contrast, when AzadC was added twice (after 0 h and additionally after 24 h), we detected an additional increase of genomic AzadC after 48 h compared to the 24 h timepoint. These results suggested that 24 h after addition, all available AzadC had been either incorporated or inactivated by hydrolysis or removal from the genome. However, taken into account that the proliferation rates of the wt and the DNMT-TKO cells were almost precisely 24 h when exposed to 2.5 µM of AzadC, these results implied that removal of genomically incorporated AzadC mostly depended on the global scale on passive dilution by cell division (Figure 2E), whereas removal by repair mechanisms or spontaneous decay only played a minor role. In consequence, to avoid passive dilution and to increase the amount of genomically incorporated AzadC, the compound has to be added continuously (Figure 2F).

**Figure 2:**
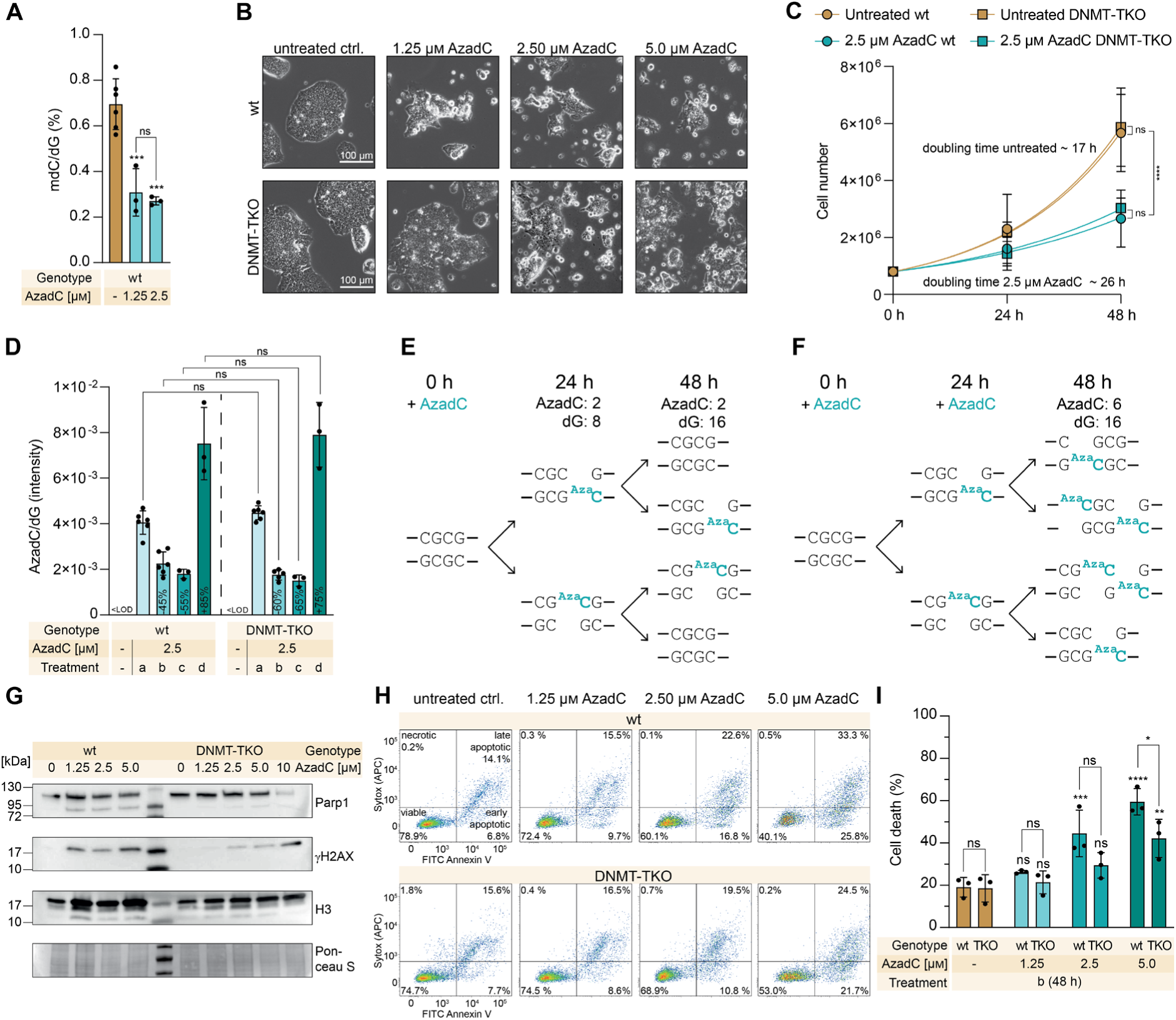
Effects of AzadC treatment in wt and DNMT-TKO mESCs under 2iL conditions. A) Amount of mdC, quantified by QQQ-MS and normalized to the amount of dG, in the wt after 48 h treatment with AzadC compared to the untreated control. Bar represents mean, error bars represent standard deviation (SD), each dot represents one biologically independent replicate. One-way ANOVA combined with Tukey’s multiple comparisons test (Supplementary Data File 1, Figure 2A). Stars above bars of AzadC-treated samples indicate significant difference of the mean compared to the untreated control. B) Representative brightfield microscopy images of wt and DNMT-TKO cells after treatment with increasing concentrations of AzadC. C) Proliferation curve of wt and DNMT-TKO cells after treatment with 2.5 µM of AzadC compared to the untreated controls. For each sample to be measured, 800,000 cells were seeded initially (0 h). For the 24 h and the 48 h timepoints, three independent biological replicates were quantified. Symbol represents mean, error bar represents standard deviation. Fitting of growth curve by exponential (Malthusian) growth with the constrain Y_0_ = 800,000 (Supplementary Data File 1, Figure 2C). D) Intensity of AzadC signal normalized to the intensity of dG signal in genomic DNA, measured by QQQ-MS, in the wt and the DNMT-TKO after treatment with 2.5 µM of AzadC. LOD = limit of detection, treatment a = AzadC addition at 0 h, harvest after 24 h; b = AzadC addition at 0 h, harvest after 48 h; c = AzadC addition at 0 h, medium change after 24 h to medium without AzadC, harvest after 48 h; d = AzadC addition at 0 h, medium change after 24 γH2AX levels in wt and DNMT-TKO cells treated with increasing concentrations of AzadC (treatment b). Histone H3 and Ponceau S staining served as a loading control. H) Representative flow cytometry scatter plots of wt and DNMT-TKO cells after 48 h treatment (treatment b) with increasing concentrations of AzadC compared to the untreated control (n = 10,000 events per condition) using FITC-Annexin V-binding as a marker for apoptosis and Sytox Red Cell Death Stain as a marker for dead cells. Viable cells are Annexin V^-^/Sytox^-^, cells in an early apoptotic state are Annexin V^+^/Sytox^-^, cells in a late apoptotic state are Annexin V^+^/Sytox^+^ and cells that have been died from other reasons (necrosis, etc.) are Annexin V^-^/Sytox^+^. I) Summary of cell death events (necrotic + early apoptotic + late apoptotic as indicated by panel H) in wt and DNMT-TKO cells after 48 h treatment (treatment b) with AzadC in increasing concentrations compared to the untreated control. Bars represent mean, error bars represent standard deviation, dots represent biologically independent replicates. TStars above bars of AzadC-treated samples indicate significant difference of the mean compared to the respective untreated control. A), D) and I) ns p_adj_ > 0.05, *0.05 > p_adj_ > 0.01, ** 0.01 > p_adj_ > 0.001, *** 0.001 > p_adj_ > 0.0001, **** p_adj_ < 0.0001.

To investigate the formation of DNA damage in the wt and the DNMT-TKO after AzadC treatment in a dose-dependent manner, we performed an immunoblot analysis against γH2AX (Figure 2G, Figure S2B). H2AX is a histone variant which is placed as a mark at sites of DNA double strand breaks (DSB) and is subsequently phosphorylated at Ser-139 (gH2AX) to recruit the repair machinery. Since one γH2AX is placed per DSB, it is a very sensitive and quantitative marker for DSB formation (45). As expected, we detected only a very faint γH2AX signal in the untreated controls of both genotypes, whereas AzadC treatment with all concentrations tested (1.25 µM – 5 µM) resulted in comparable formation of DSBs in the wt. In the DNMT-TKO, we observed a strict dose-dependent formation of DSBs with no detectable signal after 1.25 µM AzadC treatment, but a clearly visible signal at the higher concentrations. In parallel, we detected the levels of Parp1, which serve not only as a marker for DNA-damage but also induction of apoptosis. It can be cleaved in a 24 kDa fragment, which remains at the lesion, and a 89 kDa fragment that is released in the cytosol to induce apoptosis (46). The total Parp1 levels did not differ between the wt and the DNMT-TKO, but with slightly lower levels in the untreated wt (Figure 2G, Figure S2B). In contrast, the 89 kDa cleaved fragment showed only a strong signal in the wt, already at a concentration of 1.25 µM of AzadC, but not in the DNMT-TKO at any concentration tested. Last, we quantified the amount of dead cells by flow cytometry after 48 h treatment with AzadC in a concentration-dependent manner. AzadC-treatment resulted in increased amounts of apoptotic, but not necrotic cells (Figure 2H) and we observed, as indicated by the brightfield microscopy images (Figure 2B), a strictly dose-dependent increase of dead cells in both genotypes (Figure 2I). Although the observed hypomethylating effect was already at a maximum in the wt with 1.25 µM of AzadC (Figure 2A) and at the same time, the level of γH2AX was already substantially increased at this concentration (Figure 2G), 1.25 µM of AzadC were not enough to significantly induce apoptosis, which is consistent with our previously reported observations in AzadC or AzaC treated human leukemic cells (44). Compared to the wt, the DNMT-TKO required higher AzadC concentrations to go into apoptosis, but the dose-depending effect was comparable (Figure 2I).

In summary, our results show that genomically incorporated AzadC is mainly removed by passive dilution during cell division unless new substance is provided at least every 24 h. AzadC already exhibits an intrinsic anti-proliferative and apoptotic effect independent from DNMT-crosslinking, but dependent on active DNA-replication. After 24 h, when AzadC concentration was at a maximum (Figure 2D), the cells had undergone one replication cycle and already significant reduction of mdC (12), apoptosis was only induced to a very low extent (Figure S2C). This result was in accordance with previous studies showing that the cytotoxic potential of AzadC unravels during the second DNA-replication cycle after incorporation (25).

### When DNMT-activity is high, DNA-DNMT crosslinking dominates the efficacy profile of AzadC

Upon priming, wt mESCs undergo reprogramming, including changing and overall increasing DNA methylation patterns (29,30) as a result of higher expression of all three DNMT enzymes and higher DNMT activity (Figure 1E, F). Therefore, we expected increased sensitivity of the wt towards AzadC treatment. Whereas AzadC-induced DNA lesions that do not originate from DNA-DNMT crosslinking are strictly dose-dependent as they only depend on the amount of incorporated AzadC, DNA lesions from DNA-DNMT crosslinking depend on the amount of incorporated AzadC but even more on the activity of DNMT enzymes. With increasing amounts of these crosslink, genome instability is multiplied. Consequently, when DNMT activity is high, a lower amount of genomically incorporated AzadC can nevertheless lead to more devastating effects than a substantially higher amount of AzadC under conditions of low DNMT activity. In line with these preliminary considerations, we did not observe any proliferation of AzadC-treated wt cells under Lif conditions (Figure S3A). Furthermore, brightfield images indicated that under Lif conditions, massive cell death already occurred in the wt when the cells were treated with as little as 0.1 µM of AzadC (Figure 3A). The drastic increase of dead cells after low-dose AzadC treatment was additionally confirmed by the flow cytometry-based apoptosis and cell death assay (Figure 3B). In contrast, the DNMT-TKO did not respond equally sensitive (Figure 3C, Figure S3A) but the primed DNMT-TKO mESCs were more sensitive towards low concentrations of AzadC (Figure 3D) than the ones under naïve conditions, which did not respond to a AzadC concentration below 2.5 µM (Figure 2B). Overall, these results showed that the DNMT-dependent effect are highly dominant for the toxicity profile of AzadC when DNMT activity is high and DNMT-independent DNA lesions only play a subordinate role (Figure 3E).

**Figure 3:**
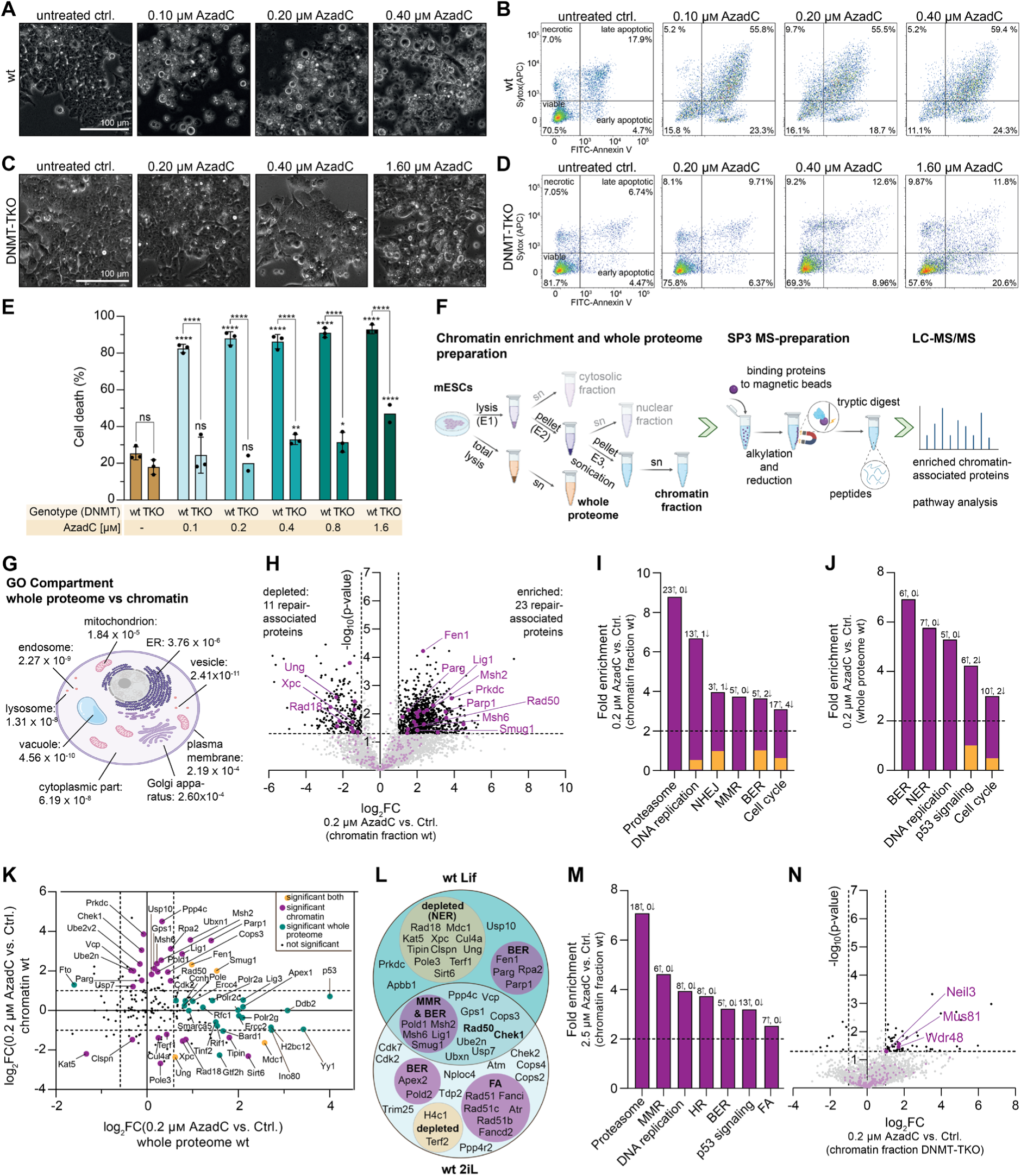
Effects of AzadC-treatment in mESCs under Lif conditions and involved DNA repair pathways under Lif and 2iL conditions. Lif conditions are 48 h pre-incubation in Lif medium, followed by 48 h treatment in Lif medium (96 h Lif in total). A) Representative brightfield microscopy images of wt cells after treatment with increasing concentrations of AzadC (Lif). B) Representative flow cytometry scatter plots of wt cells (Lif) with increasing concentrations of AzadC compared to the untreated control (n = 10,000 events per condition) using FITC-Annexin V-binding as a marker for apoptosis and Sytox Red as a marker for dead cells. C) Representative brightfield microscopy images of DNMT-TKO cells after treatment with increasing concentrations of AzadC (Lif). D) Representative flow cytometry scatter plots of DNMT-TKO cells (Lif) with increasing concentrations of AzadC compared to the untreated control (n = 10,000 events per condition). E) Summary of cell death events (necrotic + early apoptotic + late apoptotic as indicated by panels B, D) in wt and DNMT-TKO cells after 48 h treatment with AzadC in increasing concentrations compared to the untreated control. Bars represent mean, error bars represent standard deviation, dots represent biologically independent replicates. Stars above bars of AzadC-treated samples indicate significant difference of the mean compared to the respective untreated control. ns p_adj_ > 0.05, * 0.05 > p_adj_ > 0.01, ** 0.01 > p_adj_ > 0.001, *** 0.001 > p_adj_ > 0.0001, **** p_adj_ < 0.0001. F) Workflow for chromatin enrichment and whole proteome analysis and subsequent LC-MS/MS measurement. E1 – E3 are different buffer formulations that can solubilize different subcellular compartments. G) Significantly enriched Gene Ontology terms (GO Compartment) when the proteins detected in the whole proteome of wt cells was compared to the respective chromatin fraction (Lif conditions). Figure panels F, G) were created with the help of BioRender.com. H) Volcano plot of chromatin-enriched proteins: after AzadC-treatment of wt cells under Lif conditions (left side: untreated control, right side: 0.2 µM AzadC-treated mESCs). Not significantly enriched proteins (-log(p-value) < 1.3 and |log_2_FC| < 1) are marked grey. Significantly enriched proteins in one of the two conditions are labelled in black. Proteins that are involved in DNA-repair according to reactome are labelled in purple. (I) Pathway enrichment analysis of the data in H) performed with pathfindR. J) Pathway enrichment analysis of the whole proteome data of AzadC-treated mESCs under Lif conditions compared to the untreated ctrl. using pathfindR. I, J) The number of enriched (I)/upregulated (J) (↑, purple) and depleted (I)/downregulated (J) (↓, orange) proteins after AzadC treatment that are assigned to the different pathways are indicated. K) Correlation plot of the expression level changes (X-axis, whole proteome) or of the chromatin enrichment changes (Y-axis, chromatin fraction) of DNA repair-associated proteins of AzadC-treated mESCs under Lif conditions in comparison to the untreated control. L) Venn diagram of significantly chromatin enriched and depleted (marked with orange circle) proteins after AzadC treatment of wt cells under Lif and 2iL conditions. Proteins that are assigned to a specific DNA repair pathway are grouped. M) Pathway enrichment analysis of the chromatin enrichment data of AzadC-treated mESCs under 2iL conditions compared to the untreated ctrl. using pathfindR. N) Volcano plot of chromatin-enriched proteins of 0.2 µM AzadC-treated mESCs DNMT-TKO under Lif conditions compared to the untreated control (left side: untreated control, right side: AzadC-treated mESCs DNMT-TKO).

### The activation of specific DNA repair pathways as a response to AzadC-induced DNA lesions depends on DNMT-activity

To investigate the DDR towards AzadC in a systematic and holistic way, we treated wt and DNMT-TKO cells with AzadC under Lif and 2iL conditions. Afterwards, we divided the cells in two portions – one to enrich the chromatin fraction to check for recruitment of DNA repair-associated proteins to the site of action, and one to isolate the whole proteome to check for their expression level changes. Subsequently, the isolated proteins were subjected to an established SP3 workflow and analyzed by LC-MS/MS (Figure 3F). To validate our workflow, we first analyzed the detected proteins in the chromatin fraction against the whole proteome and vice versa. GO term analysis revealed that in the chromatin fraction, GO-Terms associated with regulation of (DNA-templated) transcription (e.g. GO:0006355, GO:0006357), DNA-binding (e.g. GO:0003677, GO:0043565, GO:0000976) and DNA-binding transcription factor activities (e.g. GO:0003700, GO:0000981, GO:0001228, GO:0001227) were highly overrepresented compared to the whole proteome (Figure S3B, Supplementary Data File 3). On the other hand, the whole proteome showed a massive enrichment for all cellular compartments except the nucleus when compared to the proteins of the chromatin fraction (Figure 3G). The comparison in both directions confirmed that the prepared chromatin-associated proteome was as anticipated an overrepresentation of nuclear and chromatin-bound proteins in comparison to the total cellular proteome.

Next, we investigated the differences of the chromatin-associated proteome between the untreated and the AzadC-treated wt cells under Lif conditions with a focus on enrichment-changes of proteins that are directly involved in DNA-repair as assigned by the reactome pathway knowledge base (35). Upon treatment, we observed an overall massive change of the chromatin-associated proteins. Among the significantly deregulated chromatin-associated proteins, 23 DNA repair-associated proteins were enriched after AzadC treatment and 11 DNA repair-associated proteins were depleted compared to the chromatin fraction of the untreated control (Figure 3H, Supplementary Data File 4 Table S4). A highly enriched DNA repair-associated protein was Parp1, which was already reported to be involved in the repair of AzadC-induced DNA lesions via base-excision repair (BER) (26) and now confirmed by our complementary approach. Parp1 catalyses the poly-ADP-ribosylation (PAR chains) of itself and several other targets, among them histones. Its catalytic activity is stimulated by DNA damage, among other factors and subsequently, the PAR chains recruit many other proteins involved in several DNA repair pathways to the DNA damage sites (47). For efficient DNA repair, the immediate turnover of the PAR chains is equally important, which is mediated by several glycohydrolases, including Poly(ADP-ribose) glycohydrolase (Parg), an enzyme that we also found to be significantly enriched (Figure 3H). Another top enriched hit was the flap endonuclease 1 (Fen1) that belongs to the structure-specific endonucleases (48). Fen1 is important for the maturation of Okazaki fragments during DNA replication but it is also required to deal with DNA replication stress caused by a stalled replication fork and involved in long-patch BER (49) and (microhomology-mediated) alternative end joining (a-EJ) (50,51). A-EJ is considered as an alternative non-homologous end joining (NHEJ) pathway that results in deletions. Therefore, this error-prone repair pathway is a mutagenic driver, but it is activated when NHEJ and homologous recombination (HR) repair pathways are compromised (52). Previous reports indicated that a-EJ depends on double-strand repair protein Mre11, Nibrin (Nbs1), Parp1, DNA repair protein Xrcc1, Fen1, Ligase I (Lig1) and Ligase III (Lig3) (51,53). In our chromatin dataset, Mre11 showed an three times fold enrichment (log_2_FC 1.65) after AzadC treatment of wt mESCs under Lif conditions but failed to reach overall the significant threshold (-log(p-value) > 1.3 and log_2_FC >|1|). However, we found Parp1, Fen1 and Lig1 being highly enriched after AzadC treatment, suggesting that a-EJ is one of the involved repair mechanisms to handle AzadC-induced DNA lesions.

Other important repair proteins that we found significantly enriched in the chromatin fraction of AzadC-treated wt cells under Lif conditions were the DNA-dependent protein kinase catalytic subunit Prkdc, the DNA mismatch repair proteins Msh2 and Msh6, the single-strand selective monofunctional uracil DNA glycosylase Smug1 and the DNA repair protein Rad50. Prkdc regulates phosphorylation of H2AX and is one of the key proteins in NHEJ (54,55). Msh2 and Msh6 are central components of the post-replicative DNA mismatch repair system (MMR) by forming the heterodimer MutSα that binds to DNA mismatches to initiate repair (56). Rad50 on the other hand plays a pivotal role in DSB repair (57). Interestingly, the uracil DNA glycosylase Ung, which belongs to the same protein superfamily as Smug1, was significantly depleted from the chromatin fraction of AzadC-treated wt cells (Figure 3H). Smug1 and Ung are both part of the BER machinery to repair the presence of uracil in the DNA after deamination of cytosine. However, while Ung is important for repair of uracil in replicating DNA, Smug1 is more important to remove uracil in non-replicating chromatin (58). Among the depleted proteins were also the E3 ubiquitin-protein ligase Rad18, which is an essential component to link DNA damage signalling to activation of HR (59), and the DNA repair protein complementing XP-C cells homolog Xpc, which is an integral component of the nucleotide-excision repair machinery (NER) (60).

To obtain a holistic picture of the affected cellular pathways, with a focus on the involved DNA repair pathways, we performed a pathway enrichment analysis using pathfindR (43) (Figure 3I, Supplementary Data File 4 Table S5). One of the top enriched terms was “Proteasome” with a more than eightfold enrichment and 23 contributing proteins being highly enriched in the chromatin fraction of the AzadC-treated wt cells compared to untreated cells (Table 5). Protein-DNA crosslinks (PDCs) can be repaired by proteasomal activity that digests the bulky protein adducts to the peptide level to allow the canonical repair machinery to excise the affected DNA sites in a next step (61,62). Our chromatin proteomics data strongly indicate that the proteasome removes the covalently trapped DNMTs, whereas other proteases involved in removal of PDCs like Spartan could not be detected in the chromatin fraction. As expected, pathway enrichment analysis also revealed that “DNA replication” and “Cell cycle” were heavily affected after AzadC-treatment as many checkpoint proteins were significantly enriched in the chromatin fraction after AzadC treatment. Among them was the serine/threonine-protein kinase Chek1, which is central for checkpoint-mediated cell cycle arrest and the activation of the DDR when DNA damage is present (63), as well as all components of the mini-chromosome maintenance (MCM) complex (Mcm2 – Mcm7) (Table 5). The MCM complex was first discovered to be required for controlled initiation and elongation of DNA replication, but since the MCM components are much more abundant in the cell than the amount of origins of replications and replication forks per cell per replication cycle would suggest, additional roles of this proteins were assumed (64). More recent studies indicated that the MCM complex also plays an important role in sensing DNA damage at replication forks and subsequently recruiting the DNA repair machinery (65,66). Our results confirmed these previous observations. Although untreated wt cells undergo active DNA replication in contrast to the AzadC-treated cells under Lif conditions (Figure S3A), we found all six components of the MCM (Mcm2 – Mcm7) highly enriched in the chromatin fraction of the AzadC-treated cells compared to the untreated cells. This emphasizes the central role of the MCM complex components for the DDR. Regarding the activated DNA repair pathways, NHEJ, MMR and BER showed strong enrichment in the pathway analysis in AzadC-treated wt cells compared to untreated cells under Lif conditions (Figure 3I and Table 5). This is in line with the upregulated chromatin-recruitment of key components of those pathways (Figure 3H), indicating that those repair pathways were activated.

**Table 5:**
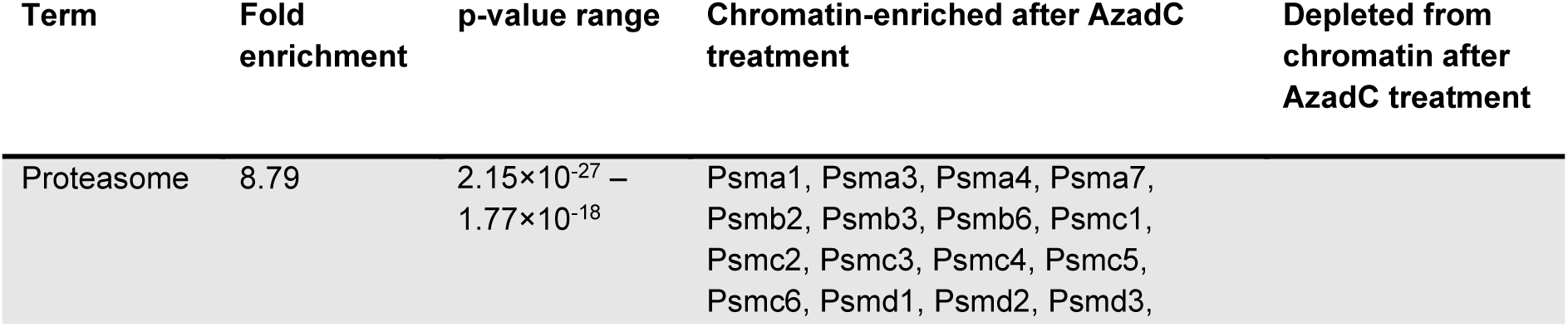
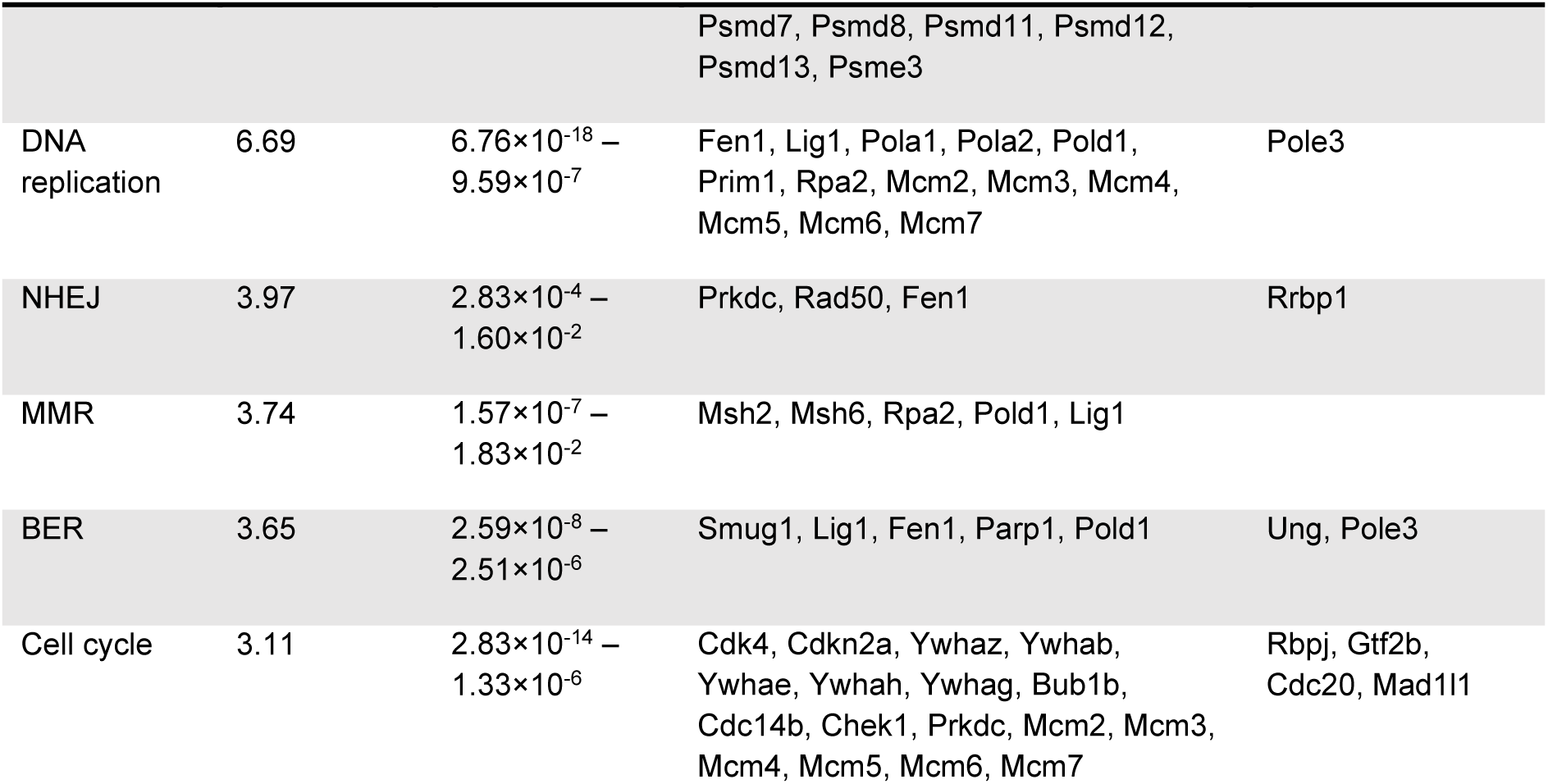
List of selected significantly enriched pathways after 0.2 µM AzadC treatment over 48 h of wt cells under Lif conditions after having analysed the chromatin-associated proteome. The proteins that are assigned to the specific pathway according to pathfindR and showed a significant change in chromatin-enrichment after treatment are listed .

To gain better insights in the activation of the DDR in wt cells under Lif conditions, we then analysed the global protein expression level changes after AzadC treatment (Supplementary Data File 4 Table S6) and subsequently compared the chromatin enrichment of the DNA repair associated proteins with their global expression level changes (Supplementary Data File 4 Table S7). Pathway enrichment analysis of the whole proteome data revealed that BER also showed a significant enrichment (Figure 3J) (Supplementary Data File 4 Table S8). Among the BER-associated proteins, Smug1 and Fen1 showed both significantly increased expression levels and significantly changed chromatin enrichment (Figure 3K). These results suggested that BER is activated by increased protein expression levels as well as by active recruitment of the involved DNA repair proteins to the site of action. Strikingly, all other DNA repair associated proteins that were found to be enriched in the chromatin fraction after AzadC-treatment did not show any significant expression level changes (Figure 3K), revealing that DDR activation in response to AzadC-induced DNA lesions is substantially based on active chromatin recruitment. On the other hand, there were several DNA repair-associated proteins that showed exclusively higher expression levels after AzadC treatment but were not significantly enriched in the chromatin fraction. The most prominent one, with a sixteen fold higher protein level after AzadC treatment (-log(p-value) = 1.4, log_2_FC = 4), was the key tumour suppressor p53. P53 is a nuclear transcription factor that plays, among many other functions, a pivotal role in DDR by inducing the expression of many DNA repair associated genes and genes that trigger cell-cycle arrest and apoptosis in the presence of DNA damage (67). Under normal conditions, p53 has a very short half-life time and is additionally mostly present in an inactivated state. Our whole proteome data are in perfect accordance with literature that upon DNA damage sensing, the p53 protein levels are substantially increased as the *Tp53* mRNA is more efficiently translated and the p53 protein is stabilized, leading to its accumulation (68). The activation of p53 and the subsequent effects were also observed in the pathway enrichment analysis that showed p53 signalling among the top enriched pathways (Figure 3J, Supplementary Data File 4 Table S8). The reason why we did not observe a significant enrichment of p53 in the chromatin fraction of AzadC-treated cells might be that transcription factors usually bind very transiently to the DNA and can therefore often not be captured without crosslinking. One of the target genes of p53 is the DNA damage-binding protein 2 (Ddb2), a key component of NER. Ddb2 recognizes UV-induced DNA lesions. It subsequently forms a complex at the lesion site with the ubiquitin ligase Cullin-4a (Cul4a), which then initiates the proteolytic digest of Ddb2. This targeted degradation at the lesion site in turn recruits the DNA repair protein Xpc to initiate global genome NER (GG-NER) (69,70). Strikingly, Ddb2 showed a significant increase in global protein levels, but we failed to detect it in the chromatin fraction, and Cul4a and Xpc were even depleted from the chromatin fraction after AzadC treatment (Figure 3H, Supplementary Data File 4 Table S4). Having a closer look on other components of the GG-NER, we observed that the majority of the depleted proteins in the chromatin fraction after AzadC-treatment belong or are at least associated to this repair pathway. Among the depleted proteins were TIMELESS-interacting protein Tipin and Claspin (Clspn), which are part of the intra-S checkpoint that is activated upon DNA replication stress (71), the NAD^+^-dependent protein deacetylase sirtuin-6 (Sirt6) and the DNA polymerase e (Pole). Clspn and Sirt6 were shown to be involved in regulation of NER via interaction with Ddb2 (72,73). Pole is in addition to DNA polymerase d (Pold) one of the DNA polymerases that is widely used to resynthesize the removed parts in different DNA repair pathways, including NER (74). Moreover, there were several NER components that were not enriched in the chromatin fraction after AzadC-treatment (Figure 3K), but significantly upregulated on the global protein expression level and consequently, pathway enrichment analysis also revealed an upregulation of NER based on the whole protein expression data (Figure 3J, K). Replication factor C (Rfc1) recruits Pold to the sites of NER (75). Ercc2 (Xpd) and Ercc4 (Xpf) are both central parts of NER. Errc2 acts as a DNA helicase and Ercc4 as the catalytic subunit of a structure-specific DNA repair endonuclease for the 5’ incision during DNA repair (76,77). Errc2 is part of the large multi-subunit transcription factor IIH (TFIIH) around which the whole NER machinery is built because it unwinds the DNA strands for repair, scans for the lesion and mediates its removal (78). Other components of the TFIIH, which we also found to be significantly upregulated in the whole proteome after AzadC-treatment but not enriched in the chromatin fraction, were Cyclin-H (Ccnh) and the general transcription factor IIH subunit 4 (Gtf2H4). These combined results suggest that when replication stress is very high after AzadC-treatment, NER is initially activated on the protein expression level but then either not used or even actively excluded from the AzadC-induced DNA lesion repair as it progresses.

As DNMT-activity and therefore the induced replication stress upon AzadC-treatment is very different in the wt under Lif and 2iL conditions, we replicated the chromatin enrichment workflow in wt cells that were treated with 2.5 µM of AzadC under 2iL conditions, with the respective untreated 2iL wt cells as a control. Under 2iL conditions, we observed 33 DNA repair-associated proteins being enriched in the chromatin fraction after AzadC treatment and only two repair-associated proteins being depleted (Figure S3C, Supplementary Data File 4, Table S9). 14 DNA repair-associated proteins were enriched in the chromatin fraction under 2iL and Lif conditions (Figure 3L). Among them were several components of BER and MMR, which were also significantly enriched as pathways under 2iL conditions (Figure 3M, Supplementary Data File 4 Table S10). Moreover, Rad50 and Chek1 were highly enriched after AzadC-treatment under both conditions, indicating that they play a central role in dealing with AzadC-induced DNA lesions. Additionally, we observed after AzadC-treatment under 2iL conditions, chromatin enrichment of the Ataxia-telangiectasia mutated (Atm) and Rad3-related (Atr) protein kinases, including their downstream targets Chek1 and Chek2 (Figure 3L, Figure S3C). Atm and Atr are essential for genome integrity, the DDR and checkpoint signalling as well as p53 activation (79,80). p53 signalling was also significantly enriched in the pathway enrichment analysis of the chromatin fraction of AzadC-treated wt cells under 2iL condition (Figure 3M). Interestingly, the FA repair pathway, including HR, that was previously reported to play an important role in repair of AzadC-induced DNA lesions (25), was under Lif conditions neither enriched as a pathway nor were the DNA repair protein Rad51 homolog 1 (Rad51, also known as Fancr) or other central components of the FA repair pathway enriched in the chromatin fraction of AzadC-treated cells. In contrast, under 2iL conditions, several important FA-proteins, including Rad51 and Fanci, were significantly enriched in the chromatin fraction after AzadC-treatment (Figure 3L, Figure S3C) and HR as well as FA, but not NHEJ were overrepresented pathways (Figure 3M). Overall, these results indicate that BER and MMR are commonly activated DDRs to deal with AzadC-induced DNA lesions. In contrast, FA and HR are only activated when DNMT-activity and therefore the resulting replication stress after DNMT-crosslinking is moderate, while NHEJ or a-EJ, which are more error prone but very efficient, are the repair pathways of choice when replication stress is immense after AzadC-treatment due to high DNMT-activity.

In a last step, we investigated which repair pathways are involved to deal with genomically incorporated AzadC when DNMT-DNA crosslinking cannot take place. To this end, we treated DNMT-TKO cells under Lif conditions with an equal dose of AzadC (200 nM over 48 h) and enriched the chromatin protein fraction. The total amount of DNA-repair associated proteins that we could detect in the chromatin fraction of DNMT-TKO cells did not differ from the wt (Supplementary Data File 4 Table S11). However, in contrast to wt cells that showed a massive change of the chromatin-bound proteins, hardly any changes of chromatin-associated proteins in general could be observed in the DNMT-TKO cells compared to the untreated control (Figure 3N). Among the few significantly enriched proteins were only three repair associated proteins: the endonuclease 8-like 3 DNA glycosylase Neil3, the crossover junction endonuclease MUS81and the WD repeat-containing protein 48 (Wdr48), which is part of the FA repair pathway. To rule out that only the low concentration of AzadC failed to activate DDR in DNMT-TKO cells, we treated the DNMT-TKO cells under 2iL conditions with 2.5 µM of AzadC over 48 h and checked chromatin enrichment. However, although we observed a loss of cellular fitness of DNMT-TKO cells (Figure 2B) and increased DNA double-strand breaks (Figure 2G) when exposed to this concentration, the chromatin-bound protein fraction was hardly affected and no concerted DDR was observed either (Figure S3D, Supplementary Data File 4 Table S12). These results indicated that without DNMT-crosslinking, AzadC does not invoke a strong and concerted DDR despite the formation of DNA lesions.

### The cytotoxic effect of AzaC depends on its DNA-incorporation and DNMT-activity but not on crosslinking of the RNA methyltransferase Trdmt1

To elucidate the potential RNA-dependent effect of AzaC, we treated wt and DNMT-TKO cells under 2iL and Lif conditions with increasing concentrations of AzaC, because a potential RNA effect should manifest in both genotypes and would be potentially dominant in the AzaC-treated DNMT-TKO mESCs (Figure 1G, Table 1). First, we checked the incorporation levels of AzaC into DNA as AzadC (Figure 4A) and into RNA as AzaC (Figure 4B) after 2.5 µM AzaC feeding for 48 h under 2iL conditions. For the wt, we observed the expected lower incorporation rate into DNA with ∼12.5% AzadC/G intensity compared to feeding with AzadC after 24 h (Figure 4A, wt treatment a). This result is in perfect accordance with literature where it was reported that 10 – 20% of AzaC are incorporated into DNA (12). The pattern that the DNA-incorporated levels dropped by half 48 h after treatment when no additional compound was added, but could be increased by additional 80% when new compound was added after 24 h, was also observed for AzaC treatment in the wt. However, while we did not detect any difference regarding the uptake and metabolization of AzadC between wt and DNMT-TKO cells (Figure 2D), AzaC incorporation into DNA was significantly lower in the DNMT-TKO compared to the wt (Figure 4A). On the other hand, the incorporation rates of AzaC into RNA were comparable between the wt and the DNMT-TKO cells (Figure 4B). Therefore, different intercellular metabolization kinetics and not a difference in the uptake of AzaC was probably the reason for the lower DNA-incorporation in the DNMT-TKO. Moreover, the level of AzaC in RNA in wt and DNMT-TKO mESCs could only be maintained but not further increased when additional compound was added after 24 h (Figure 4B). This result indicates that RNA compared to DNA has a higher turnover and AzaC could therefore not accumulate. Next, we investigated the phenotypic changes after AzaC-treatment (2.5 µM for 48 h, treatment b) in wt and DNMT-TKO cells under 2iL conditions. Whereas the proliferation rate initially slowed down in the first 24 h after AzaC treatment in the DNMT-TKO cells, proliferation quickly resumed in those cells and the cell number after 48 h was the same as for the untreated control (Figure 4C). In contrast, the wt showed a persistent slower proliferation rate after AzaC treatment with doubling times comparable to AzadC treatment. Intriguingly, the slower proliferation rate in the wt was not reflected in a higher cytotoxicity as indicated by brightfield microscopy (Figure 4D) and the flow cytometry-based apoptosis assay (Figure 4E). In the DNMT-TKO, AzaC did not have any cytotoxic activity in the applied concentration either (Figure 4D, E). When we had a closer look at the DNA hypomethylating effect of AzaC compared to AzadC in the wt cells under 2iL conditions, we observed a substantial decrease of mdC for both compounds with no difference between AzadC and AzaC at 1.25 µM and 2.5 µM (Figure 4F). In contrast, γH2AX levels in the wt were much higher after AzadC compared to AzaC treatment as indicated by the intensity of the γH2AX signal in fluorescence microscopy (Figure 4G). When we checked the chromatin-enriched proteome of wt cells after AzaC-treatment (Supplementary Data File 4, Table S13), we observed a very high correlation compared to the AzadC-treated cells (Pearson r = 0.7778, Supplementary Data File 4 Table S14) but only few repair-associated proteins reached the significance threshold (Figure 4H). In summary, those results suggest that under steady conditions, the RNA- incorporation of AzaC leads to an initial proliferation delay that can be, however, quickly overcome by the cells due to high RNA turnover. As a consequence, no persistent RNA-dependent effect can be achieved without additional AzaC supply. As DNA damage seemed to be marginal under 2iL conditions after AzaC treatment in the wt, which is in line with the absent toxicity, the inhibition of DNMTs and the resulting significant decrease in mdC appears to be responsible for the significantly decreased proliferation rate in the wt after AzaC-treatment. Although AzaC incorporation into gDNA reaches only 12.5% compared to AzadC treatment in the wt, the changes of the chromatin-associated proteins are highly similar for AzaC- and AzadC-treatment but less pronounced after AzaC treatment.

**Figure 4:**
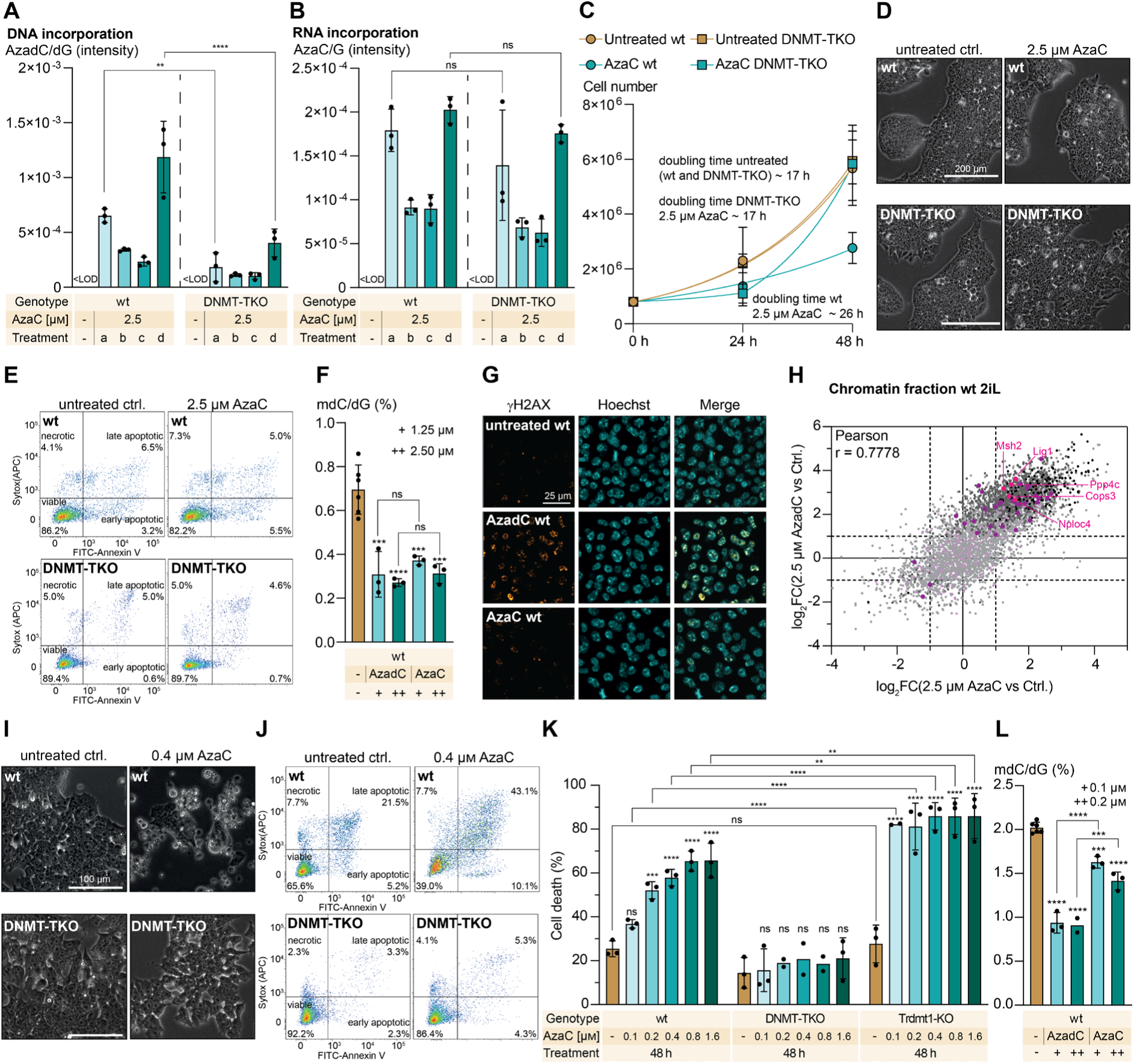
The effect of AzaC-treatment on proliferation rate and viability. A, B) Intensity of AzadC signal normalized to the intensity of dG signal in genomic DNA (A) and of AzaC in RNA (B), measured by QQQ-MS, in the wt and the DNMT-TKO after treatment with 2.5 µM of AzaC under 2iL conditions. LOD = limit of detection, treatment a = AzaC addition at 0 h, harvest after 24 h; b = AzaC addition at 0 h, harvest after 48 h; c = AzaC addition at 0 h, medium change after 24 h to medium without AzaC, harvest after 48 h; d = AzaC addition at 0 h, medium change after 24 Proliferation curve of wt and DNMT-TKO cells after treatment with 2.5 µM of AzaC under 2iL conditions compared to the untreated controls, which were also displayed in Figure 2C. For each sample to be measured, 800,000 cells were seeded initially (0 h). For the 24 h and the 48 h timepoints, three independent biological replicates were quantified. Symbol represents mean, error bar represents standard deviation. Fitting of growth curve for wt (untreated and treated) and untreated DNMT-TKO by exponential (Malthusian) growth with the constrain Y_0_ = 800,000. (Supplementary Data File 1, Figure 4C). D) Representative brightfield microscopy images of wt and DNMT-TKO cells untreated or after treatment with 2.5 µM AzaC under 2iL conditions for 48 h. E) Representative flow cytometry scatter plots of wt and DNMT-TKO cells untreated and after 48 h treatment with 2.5 µM AzaC under 2iL conditions (n = 10,000 events per condition) using FITC-Annexin V-binding as a marker for apoptosis and Sytox Red as a marker for dead cells. F) Amount of mdC, quantified by QQQ-MS and normalized to the amount of dG, in the wt under 2iL conditions after 48 h treatment with AzadC or AzaC compared to the untreated control. Ctrl. and AzadC data were also displayed in Figure 2A. Bar represents mean, error bars represent standard deviation (SD), each dot represents one biologically independent replicate. One-way ANOVA combined with Tukey’s multiple comparisons test (Supplementary Data File 1, Figure 4F). Stars above bars of AzadC or AzaC-treated samples indicate significant difference of the mean compared to the untreated control. G) Fluorescence microscopy images of γH2AX signal in wt cells (2iL conditions) untreated or after 48 h treatment with 5.0 µM AzadC or AzaC. Hoechst staining shows nuclei. H) Correlation plot of the chromatin enriched proteome changes after Aza treatment (X-axis) and AzadC treatment (Y-axis) compared to the untreated control in wt cells under 2iL conditions. Repair-associated proteins that were significantly enriched (-log(p-value) > 1.3 and |log_2_FC| >1) after both treatments are labelled magenta dots. Repair-associated proteins that were only significant after AzadC treatment are dark purple, non-significant repair associated proteins are displayed light purple. Pearson correlation is shown. Representative brightfield microscopy images of wt and DNMT-TKO cells untreated or after treatment with 0.4 µM AzaC under Lif conditions for 48 h. J) Representative flow cytometry scatter plots of wt and DNMT-TKO cells untreated and after 48 h treatment with 0.4 µM AzaC under Lif conditions (n = 10,000 events per condition) using FITC-Annexin V-binding as a marker for apoptosis and Sytox Red as a marker for dead cells. Summary of cell death events (necrotic + early apoptotic + late apoptotic) in wt, DNMT-TKO and Trdmt1-KO cells (Lif conditions) after 48 h treatment with AzaC in increasing concentrations compared to the untreated control. Bars represent mean, error bars represent standard deviation, dots represent biologically independent replicates. TStars above bars of AzaC-treated samples indicate significant difference of the mean compared to the respective untreated control. L) Amount of mdC, quantified by QQQ-MS and normalized to the amount of dG, in the wt under Lif conditions after 48 h treatment with AzadC or AzaC compared to the untreated control. Bar represents mean, error bars represent standard deviation (SD), each dot represents one biologically independent replicate. One-way ANOVA combined with Tukey’s multiple comparisons test (Supplementary Data File 1, Figure 4L). Stars above bars of AzadC or AzaC-treated samples indicate significant difference of the mean compared to the untreated control. A), B). F), K, L) ns p_adj_ > 0.05, * 0.05 > p_adj_ > 0.01, ** 0.01 > p_adj_ > 0.001, *** 0.001 > p_adj_ > 0.0001, **** p_adj_ < 0.0001.

Next, we investigated the effect of AzaC when the transcriptome and proteome and therefore cellular identity of both genotypes drastically change by switching the mESC culture conditions from naïve to primed. In stark contrast to 2iL conditions, AzaC-treatment of wt cells under Lif conditions had a very strong effect on the cellular viability as indicated by brightfield microscopy (Figure 4I) and the flowcytometry-based apoptosis assay (Figure 4J). Similar to AzadC-treatment, cellular proliferation of the wt completely stopped using an AzaC concentration as low as 0.2 µM (Figure S4A). The effect on the viability of DNMT-TKO cells, however, was minimal at this concentration (Figure 4I, J) and the proliferation rate was not affected either (Figure S4A). To further investigate the RNA-dependent features of AzaC, we quantified cell death in wt, DNMT-TKO and additionally in Trdmt1-deficient (Dnmt2-deficient) mESCs (Trdmt1-KO) after exposing the cells to increasing AzaC concentrations for 48 h under Lif conditions. In the Trdmt1-KO, no Trdmt1-crosslinking to RNA by AzaC can take place. For the wt, we observed a steady increase in toxicity with increasing concentrations of AzaC, whereas for the DNMT-TKO, AzaC was not toxic at the applied concentrations (Figure 4K). Remarkably, the Trdmt1-KO was significantly the most sensitive genotype towards AzaC-treatment, which additionally contradicts the idea that Trdmt1-RNA crosslinking as a result of RNA-incorporation of AzaC has a negative impact on cellular well-being. If Trdmt1-RNA crosslinking by AzaC had a negative impact on the proliferation rate and enforced induction of cell death analogous to the effects of DNMT-DNA-crosslinking for DNA-incorporated AzaC, the Trdmt1-KO would be less sensitive towards AzaC-treatment and not more.

Last, we wanted to check whether DNA-damage or decreased mdC levels were responsible for the substantially increased cell death events in the wt under Lif conditions. First, we quantified mdC levels in the untreated control, AzadC- and AzaC-treated cells and observed that although mdC levels were significantly decreased after AzaC-treatment compared to the untreated control, the levels remained at significantly higher levels compared to AzadC-treated cells with no difference between the two concentrations tested (Figure 4L). Under 2iL conditions, mdC levels were equally reduced after AzaC and AzadC-treatment (Figure 4F), cytotoxicity of AzaC, however, remained low. In contrast, under Lif conditions, AzaC was significantly less efficient compared to AzadC to reduce mdC (Figure 4L), but cytotoxicity was high. Therefore, we concluded that the induction of substantial DNA-damage after AzaC-treatment because of high DNMT-activity was the underlying reason for the dramatically increased sensitivity of the primed wt mESCs towards AzaC compared to the naïve wt mESCs. To substantiate this hypothesis, we additionally tested the non-nucleoside DNMT inhibitor RG-108 (31) on wt cells under Lif conditions and although we reached a reduction of mdC comparable to the reduction after AzaC-treatment (Figure S4B), we failed to detect increased cell death (Figure S4C), showing that the reduced mdC levels did not impair cellular viability under these conditions.

In summary, the effects of AzaC appear to primarily rely on its incorporation into DNA and while DNMT-inhibition and thereby reduced mdC levels significantly slow down proliferation rate, the induction of cell death depends on the severity of the induced DNA damage. At moderate DNMT activity, the level of severe DNA-lesions and therefore cell death rates remain low after AzaC treatment, whereas under high DNMT-activity, AzaC has substantial cytotoxicity comparable to AzadC.

## DISCUSSION

Our comparison of wt mESCs to DNMT-TKO mESCs revealed that the presence of the non-canonical nucleobase 5-azacytosine contributes to the toxicity profile of AzadC independent from DNA-DNMT crosslinking. However, although the presence of 5-azacytosine in the genome results in DNA-lesions with increasing concentrations, a concerted DDR is only invoked after DNA-DNMT crosslinking. Using chromatin enrichment followed by proteome analysis of the enriched fraction, we systematically analysed the involved repair mechanisms to deal with the AzadC-induced DNMT-DNA crosslinks in a holistic manner and under different levels of DNMT-activity. Under moderate and high DNMT-activity, the DNA-DNMT crosslinks are targeted by the proteasome and trigger MMR and BER. Using our approach, we confirmed the previously reported involvement of PARP1 (26) and FA-dependent HR (25) to repair AzadC-induced lesions. However, our results indicate that under high DNMT-activity, with consequently very high replication stress after DNA-DNMT crosslinking, FA-dependent repair does not contribute anymore substantially to the repair of AzadC-induced DNA lesions but is replaced by NHEJ and a-EJ. Overall, the here presented method to compare proteome information after chromatin enrichment to whole proteome data revealed highly relevant proteome changes for chromatin dynamic-relevant processes that would have remained undiscovered if only whole proteome changes or transcriptome changes would have been investigated. These results emphasize that the spatial resolution of the cellular proteome is equally important as temporal resolution of protein expression changes over time. Furthermore, we could show that at high DNMT-activity, the formation of DNA-DNMT crosslinks and the resulting DNA damage are the dominating MoA of AzadC and AzaC, even at low concentrations, whereas an RNA-dependent effect could not be observed. DNA demethylation by DNMT inhibition on the other hand, seemed to have an impact on the proliferation rate but not on the viability of the cells. Our data suggest that the better efficacy profile of AzaC in some cancers probably relies on different uptake and metabolization kinetics rather than on an RNA-based effect. Furthermore, although the presence of AzadC in the genome results in DNA-DNMT crosslinks, which trigger a strong DDR, the global removal of AzadC from the genome primarily depends on passive dilution by ongoing replication. To keep the level of genomically incorporated AzadC high, a constant exposure to AzadC has to be guaranteed as it is expected from the low stability of AzadC towards hydrolysis (81) and now showed by our time course experiments following different treatment regimens.

In this study, we used mESCs as a model because they feature an intact DDR, which was important to investigate the DNA repair mechanisms in a comprehensive way, Moreover, they allow control over DNMT activity and dynamics by changing the culturing conditions, thereby mimicking cancer types with a very different DNMT-activity profile without having completely different cellular features. In a next step, the here acquired information on the MoAs of AzadC and AzaC have to be transferred to clinically relevant models. Previous reports showed that neither somatic mutations, including those in DNA methylation-relevant genes, nor the reduction of mdC after AzadC- and AzaC treatment provide any predictive value on therapy outcome (3,44). Our data strongly suggest that biomarkers for decitabine and azacytidine should be investigated directly on the protein (activity) level and not solely on the genome level, with cancer cells that feature a high DNMT1 activity, indicated by high global mdC level, being potentially more susceptible towards AzadC and AzaC-treatment. Furthermore, our data provide a solid basis to investigate the effects of an altered DNA-repair machinery to deal with AzadC-induced DNA lesions. Importantly, the information of alterations of the DDR should be combined with information on DNMT-activity as we could show that the level of replication stress has an influence on the choice of DNA-repair pathway. Since Rad50 was strongly enriched in our model system in the chromatin fraction after AzadC-treatment under moderate and high DNMT-activity, RAD50 should be considered as clinical relevant target to increase the efficacy of AzadC-treatment.

## DATA AVAILABILITY

The mass spectrometry proteomics data have been deposited to the ProteomeXchange Consortium (82) via the PRIDE (83) partner repository with the dataset identifier PXD045353.

Other original data, analysis files and respective metadata, which are not in the supplementary files, have been deposited using figshare.com and can be accessed via:

10.6084/m9.figshare.24146604 (apoptosis assay data)

10.6084/m9.figshare.24146592 (QQQ data)

## SUPPLEMENTARY DATA

Supplementary Data is provided in different files as indicated in the main text. Supplementary Figures, including their figure legends, are displayed in the Supplementary Figure File. Information on the statistical analysis of different data is provided in the Supplementary Data File 1. Supplementary data that is relevant for the Materials and Methods section is provided in Supplementary Data File 2. Information on the GO term analysis when chromatin-associated proteome enrichment was compared to whole proteome data can be found in Supplementary Data File 3. All other information regarding the different proteomics data sets is provided in Supplementary Data File 4.

All Supplementary Data files are available online.

## AUTHOR CONTRIBUTIONS

Tina Aumer: Data curation, Formal Analysis, Investigation, Supervision, Validation, Writing – review & editing. Linda Bergmayr: Data curation, Formal Analysis, Investigation, Supervision, Validation. Stephanie Kartika: Investigation, Validation, Writing – review & editing. Theodor Zeng: Investigation. Qingyi Ge: Investigation. Grazia Giorgio: Investigation. Maike Däther: Supervision. Alexander Hess: Investigation. Stylianos Michalakis: Funding acquisition, Supervision, Writing – review & editing. Franziska Traube: Conceptualization, Data curation, Formal Analysis, Funding acquisition, Investigation, Methodology, Project administration, Supervision, Validation, Visualization, Writing – original draft.

## Supporting information

Contains all Supplementary Data except Supplementary Figures

## ACKNOWLEDGEMENTS

We thank Dr. Matthias Heiß and Kerstin Kurz for QQQ maintenance, trouble shooting and quality checks and Dr. Matthias Heiß also for his valuable input for the QQQ measurements. We thank Dr. Pavel Kielkowski, Dmytro Makarov and Andreas Wiest for Eclipse maintenance, calibration and trouble shooting. We thank the Biomedical Center Munich Core Facility Flow Cytometry (LMU) for their support. We thank Johannes Pforr (TUM Natural School of Sciences) for his help with initial experiments for this study and Corinna Pleintinger for checking the integrity of AzaC and AzadC at the HPLC.

