## Supplementary Figures for "The type of DNA damage response after Decitabine treatment depends on the level of DNMT activity"

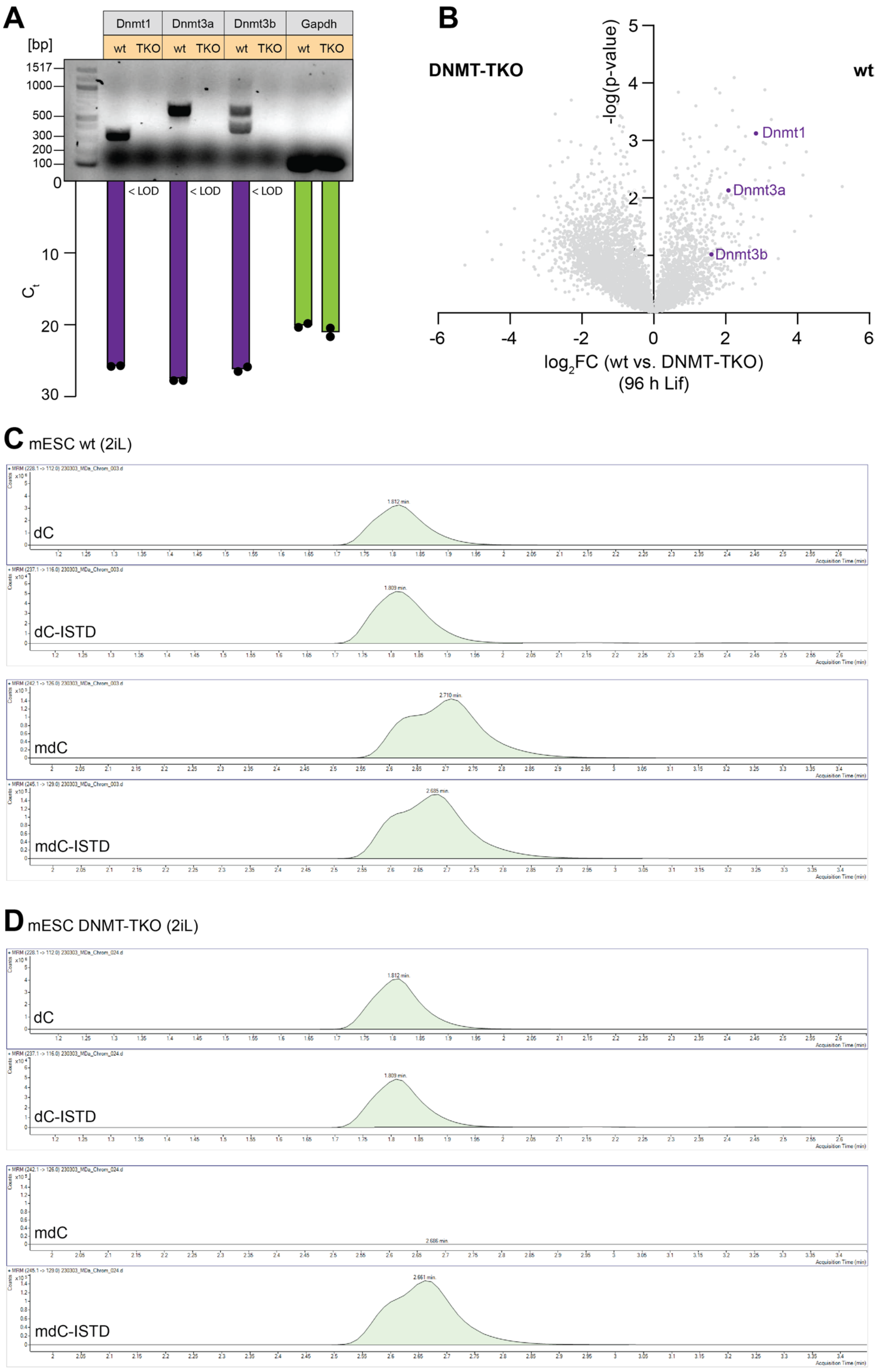


Figure S1: **Review of DNMT protein levels and activity in mESC DNMT-TKO cells.** A) RT-qPCR in mESCs wt vs DNMT-TKO (2iL conditions) using the PCR primers described in *Tsumura et al., 2006* (1) showing no signal/amplification product for Dnmt1, Dnmt3a and Dnmt3b in the DNMT-TKO compared to the wt. Gapdh served as an internal control. B) Volcano plot showing that all three DNMTs are overrepresented in the wt compared to the DNMT-TKO on the protein level. C, D) QQQ-MS chromatograms showing that there is in contrast to the wt (C) no detectable mdC in the DNMT-TKO (D), proofing the absence of any DNMT activity in the DNMT-TKO. dC and isotopologue standard (ISTD) signals of dC and mdC served as a control.

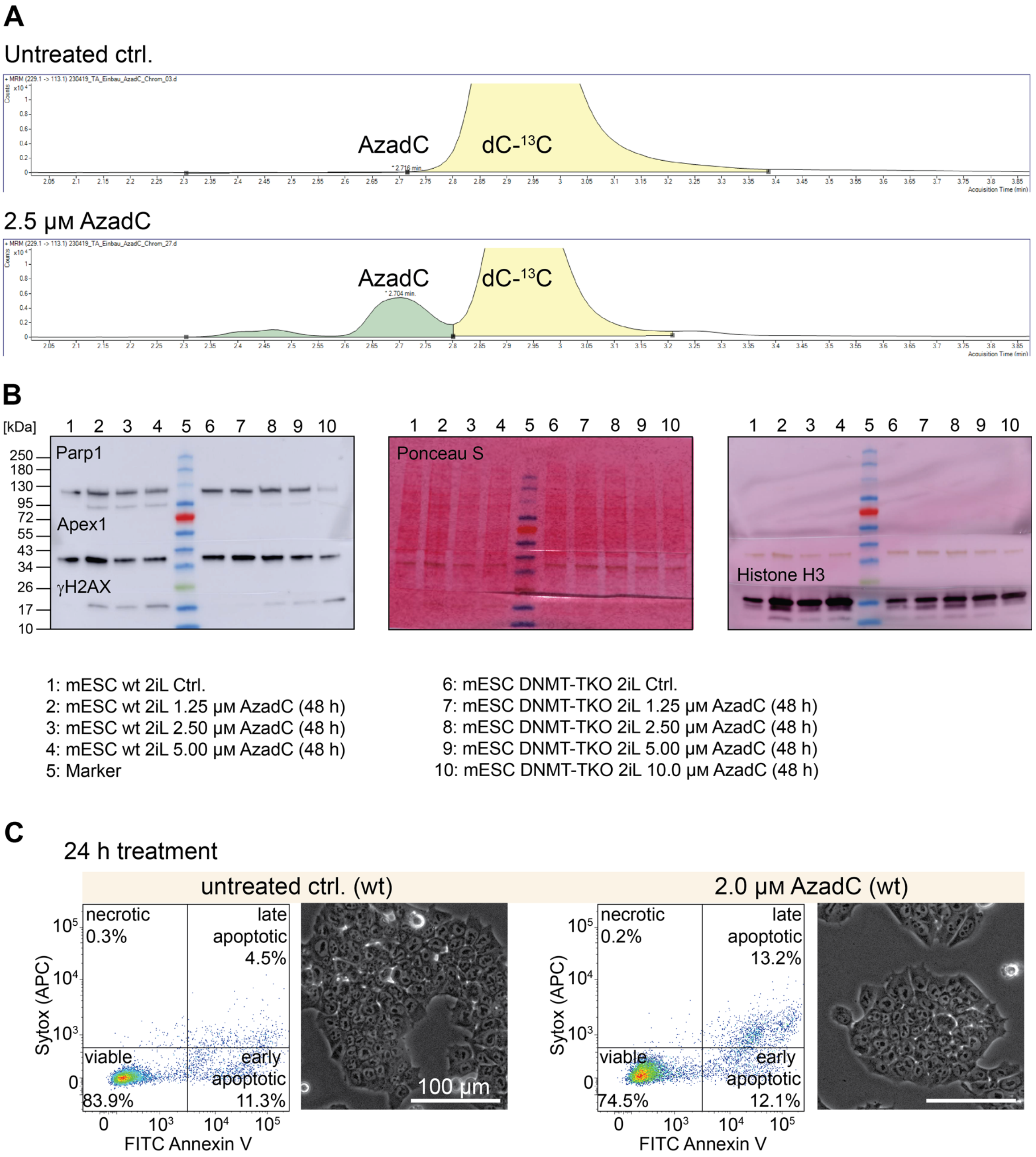


Figure S2: **Effects of AzadC treatment on wt and DNMT-TKO mESCs under 2iL conditions.** A) QQQ-MS signal of AzadC in the untreated ctrl. and after 24 h of AzadC treatment exemplarily shown for the wt. As AzadC and dC with one out of the nine carbon atoms being a $^{13}C$ (abundance ca. 1% of $^{12}C$) have a mass difference that cannot be resolved by QQQ-MS, the dC-$^{13}C$ signal is also captured when measuring AzadC. However, the chromatogram shows that both molecules can be clearly distinguished from each other by their retention time with no AzadC signal in the untreated ctrl. B) Complete images of the displayed immunoblot analysis in Figure 2G (Apex1 not shown there). In all three panels, the same blot is shown with first parallel detection of Parp1, Apex1 and γH2AX, then Ponceau S staining and afterwards stripping and then detection of Histone H3. C) Representative FACS scatter dot plots and brightfield images of wt untreated control and 2 µM AzadC-treated cells showing that after 24 h only a minor increase in dead cell events could be observed.

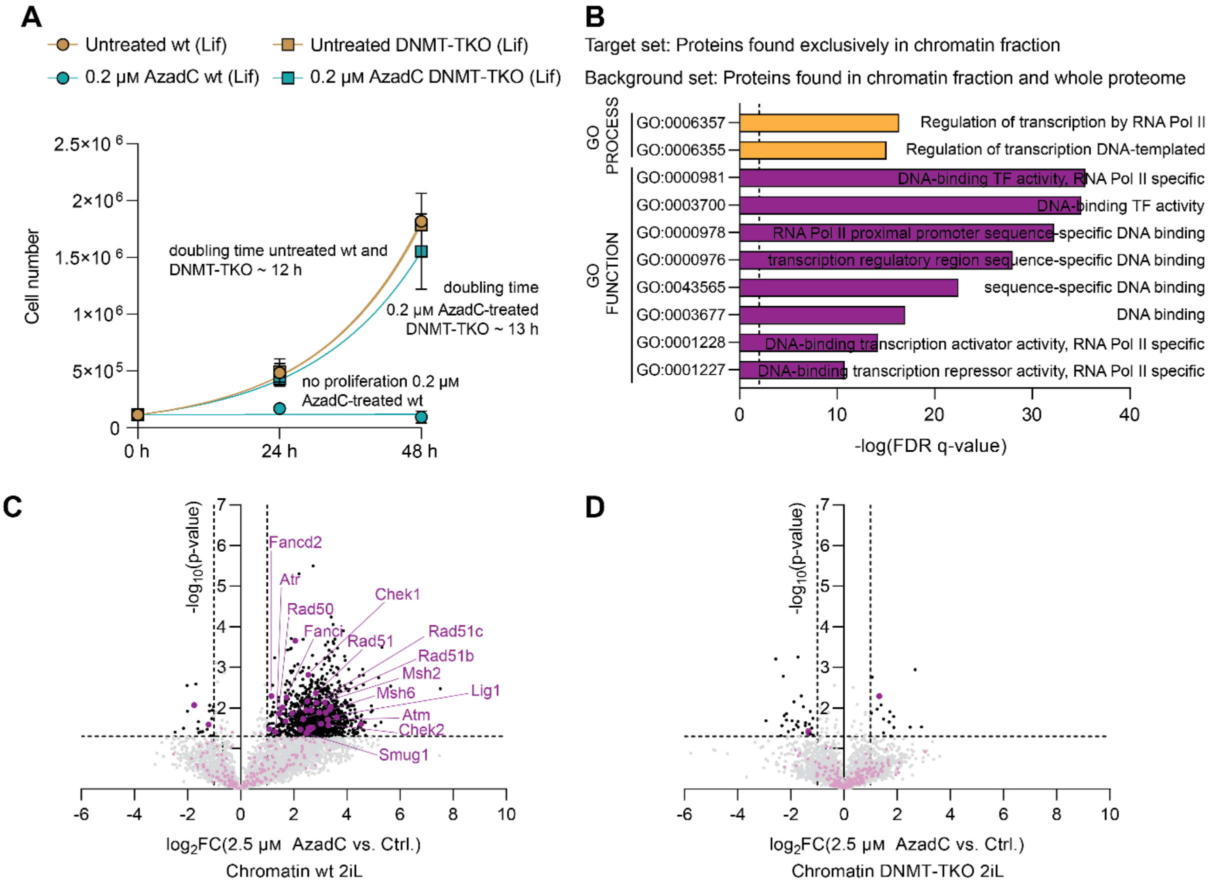


Figure S3: **Effects of AzadC-treatment in mESCs under Lif conditions and chromatin enrichment and whole proteome results of AzadC-treated mESCs compared to the untreated control.** A) Proliferation curve of wt and DNMT-TKO cells after treatment with 0.2 µM of AzadC compared to the untreated controls. For each sample to be measured, 115,000 cells were seeded initially (0 h). For the 24 h and the 48 h timepoints, three independent biological replicates were quantified. Symbol represents mean, error bar represents standard deviation. Fitting of growth curve by exponential (Malthusian) growth with the constrain Y0 = 115,000 (Supplementary Data File 1, Figure S3A).C) Volcano plot of chromatin-enriched proteins: after AzadC-treatment of wt cells under 2iL conditions (left side: untreated control, right side: 2.5 µM AzadC-treated mESCs). Not significantly enriched proteins (-log(p-value) < 1.3 and $|log_2FC| < 1$) are marked grey. Significantly enriched proteins in one of the two conditions are labelled in black. Proteins that are involved in DNA-repair according to reactome are labelled in purple. D) Volcano plot of chromatin-enriched proteins: after AzadC-treatment of DNMT-TKO cells under 2iL conditions (left side: untreated control, right side: 2.5 µM AzadC-treated mESCs). Not significantly enriched proteins (-log(p-value) < 1.3 and $|log_2FC| < 1$) are marked grey. Significantly enriched proteins in one of the two conditions are labelled in black. Proteins that are involved in DNA-repair according to reactome are labelled in purple.

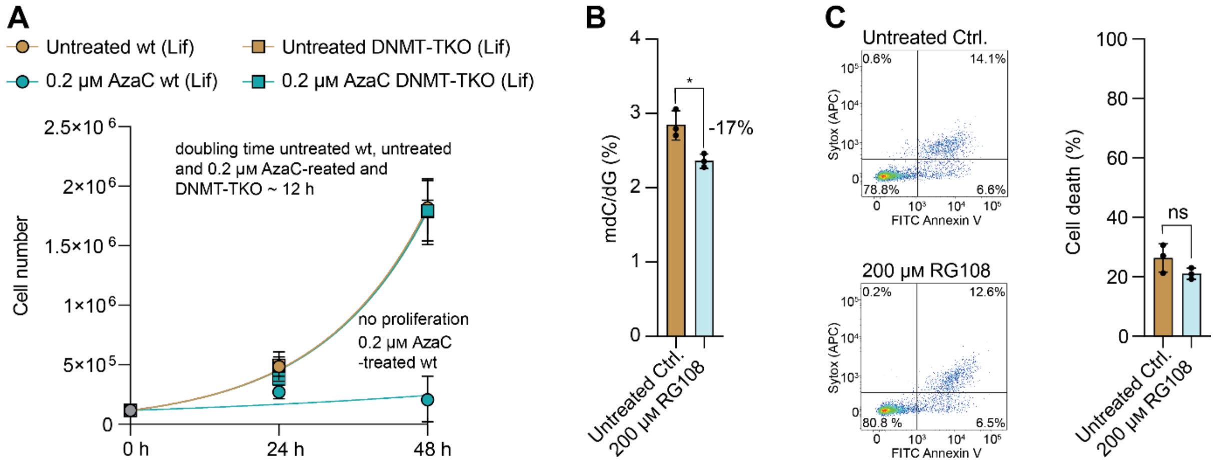


Figure S4: **Effects of AzaC and RG108 treatment in mESCs under Lif conditions.** Proliferation curve of wt and DNMT-TKO cells after treatment with 0.2 µM of AzaC under Lif conditions compared to the untreated controls, which were also displayed in Figure S3A. For each sample to be measured, 115,000 cells were seeded initially (0 h). For the 24 h and the 48 h timepoints, three independent biological replicates were quantified. Symbol represents mean, error bar represents standard deviation. Fitting of growth curve for wt (untreated and treated) and untreated DNMT-TKO by exponential (Malthusian) growth with the constrain Y0 = 115,000. (Supplementary Data File 1, Figure S4A). B) Amount of mdC, quantified by QQQ-MS and normalized to the amount of dG, in the wt after 4 weeks pre-treatment with 50 µM RG108 (2iL), followed by 48 h treatment with 50 µM RG108 and additional 48 h treatment with 200 µM RG108 under Lif conditions compared to the untreated control. Bar represents mean, error bars represent standard deviation (SD), each dot represents one biologically independent replicate. Unpaired t-test (two-sided) (Supplementary Data File 1, Figure S4B). C) Representative FACS scatter dot plots (one out of 3 n) of the flow cytometry-based apoptotic assay of wt mESCs untreated control and 200 µM RG108-treated under Lif conditions and quantification of the assay for all three biologically independent replicates. Bar represents mean, error bars represent standard deviation (SD), each dot represents one biologically independent replicate. Unpaired t-test (two-sided) (Supplementary Data File 1, Figure S4C).

### References Supplementary Figures File:

1. Tsumura, A., Hayakawa, T., Kumaki, Y., Takebayashi, S., Sakaue, M., Matsuoka, C., Shimotohno, K., Ishikawa, F., Li, E., Ueda, H.R. *et al.* (2006) Maintenance of self-renewal ability of mouse embryonic stem cells in the absence of DNA methyltransferases Dnmt1, Dnmt3a and Dnmt3b. *Genes Cells*, **11**, 805-814.
